# Expanding the Enzymatic Landscape for Polyurethane Degradation of Novel Bacterial Urethanases

**DOI:** 10.64898/2026.02.11.705263

**Authors:** Laura Rotilio, Rune R. Østergaard, Emma M. Thiesen, Pedro Paiva, Kelly Marie Dwyer, Martin B. Johansen, Andreas Sommerfeldt, Alexander Sandahl, Malene B. Keller, Suzana Siebenhaar, Rosie Graham, Daniel E. Otzen, Pedro A. Fernandes, Maria J. Ramos, Peter Westh, J. Preben Morth

## Abstract

Polyurethanes (PURs) represent a significant challenge in plastic waste management due to their chemical resilience and limited recycling options. In this study, we report the identification and characterization of five novel bacterial urethanases, expanding the enzymatic repertoire for targeted PUR depolymerization. These enzymes demonstrated carbamate-cleaving activity optimally under alkaline conditions, maintaining stability across a pH range of 7 to 10 and varying thermal and solvent tolerances. Two candidate enzymes, u17 u15 collectively exhibited high activity, catalytic efficiency, and thermostability, establishing a strong foundation for further optimization. Among them, u15 emerged as particularly notable for its catalytic efficiency on the carbamate model substrate di-urethane ethylene methylenedianiline, DUE-MDA, with a k_cat_/K_M_ of 51.8 ± 0.1 (s^-1^mM^-1^). and this motivated its selection for detailed structural analysis. High-resolution crystallography of u15 revealed key active-site architecture, including the conserved amidase signature catalytic triad and flexible loop regions that influence substrate binding and specificity. Molecular docking and molecular dynamics simulations further elucidated substrate binding determinants of u15 during urethane bond hydrolysis. Docking of DUE-MDA revealed two distinct substrate orientations (Pose A and Pose B) differing in the positioning of the carbamate group relative to Ser177. Pose A was more stable and catalytically competent, maintaining the substrate within the oxyanion hole and sustaining optimal geometry for nucleophilic attack by Ser177. Comparable behavior was observed for the partially hydrolyzed intermediate mono-urethane ethylene methylenedianiline, MUE-MDA, indicating a conserved binding mode across substrates. To further assess enzymatic performance on a realistic industrial material, the panel was then tested on a generic flexible foam substrate derived from 2,4- and 2,6-toluene diisocyanate (TDA), where u15 and u17 emerged as the most active candidates. Collectively, we benchmark the structural framework presented by enzymes in the amidase signature family as a strong foundation for further optimization aiming at advancing sustainable and scalable biocatalytic recycling of polyurethanes.

## INTRODUCTION

Synthetic polymers, such as polyurethane, nylon, and polyethylene, have become an integral part of countless industrial and consumer applications due to their low manufacturing cost, chemical versatility, and mechanical durability ^1–3^. However, these same properties contribute to persistent plastic waste problems the world is facing today ^1^.

Polyurethanes (PURs) account for about 5% of total plastic production and are valued for their versatility in flexible foams, insulation, and elastomers ^4^. Their chemical architecture makes them highly resistant to degradation, resulting in significant waste management challenges ^5^. Landfilling and incineration of PURs perpetuate environmental issues, while recycling options are limited: mechanical recycling applies only to thermoplastic PURs, and chemical recycling often requires harsh processes and leads to complex product mixtures that need extensive purification ^6–15^. Given these challenges, focus has shifted toward the development of biocatalytic depolymerization as a low energy, recycling strategy. Bioengineered enzymes, operating under mild conditions, can cleave polymers, such as Polyethylene terephthalate (PET), into reusable monomers for closed-loop production ^16–19^. The case of PET degradation, fueled by the discovery and engineering of PETase and MHETase, has highlighted how tailor-made enzymes can achieve commercial-scale recycling efficiency ^20,21^. For PURs, however, most discovered microbial hydrolases, such as lipases and cutinases, are esterases and thus primarily target the ester-based soft segments, leaving the urethane-based hard segments unaffected ^22,23^. A notable recent advancement was the identification of urethane-hydrolyzing enzymes capable of breaking down pre-treated PUR glycosylates ^24^ and methylene diphenyl diisocyanate(MDI)-based polyurethane and polyester-based PU foam ^25^. This was followed by structural and mechanistic studies that have revealed fundamental catalytic details elucidating how these enzymes function ^26^. Despite these advances, their specificity, catalytic efficiency, and thermal stability remain distant from industrial feasibility. Consequently, there is an urgent demand to discover and engineer urethanases with expanded substrate scope and enhanced catalytic performance. Similar to how PETase optimization revolutionized PET recycling, progress in urethanase discovery, characterization, and stabilization could establish a viable biocatalytic route for polyurethane recycling. Here, we report the discovery and characterization of five new bacterial urethanases identified bioinformatically, expanding beyond 60 % sequence identity from initially reported UMG-SP1, UMG-SP2, and UMG-SP3. These enzymes are amidase signature (AS) and reported urethane catalytic activity ^27,28^. Benchmarking the wide potential of the enzymatic family towards polyurethane degradation. These selected candidates were systematically assessed for catalytic activity, thermal stability, and substrate specificity against both model urethane compounds and representative PUR fragments. Among the tested candidates, urethanase 15 (u15) and urethanase 17 (u17) exhibited pronounced catalytic activity toward di-urethane ethylene Methylenedianiline (DUE-MDA, Figure S1A) ^27^, di-urethane ethylene diaminotoluene (DUE-TDA, Figure S1B) and a generic Flexible PUR foam ^29^ (Figure S1C). The crystal structure of u15 was determined by X-ray crystallography, which prompted molecular docking and molecular dynamics simulations to probe active-site architecture and rationalize sequence–function relationships. Our study integrates enzyme discovery, molecular-level mechanistic insights, and structure–function correlation, laying the groundwork for engineering next-generation biocatalysts for efficient, scalable, and sustainable PUR recycling.

## EXPERIMENTAL SECTION

### Urethanase candidate bioinformatic mining

Putative urethane-degrading enzyme candidates were identified using three query proteins, UMG-SP1 (NCBI accession: WBR49956.1), UMG-SP2 (NCBI accession: WBR49957.1), and UMG-SP3 (NCBI accession: WBR49958.1) ^24^. Homologs were retrieved with BLASTP (web) against the non-redundant (nr) and environmental non-redundant (env_nr) protein databases (accessed August 2025) ^30,31^. Hits were retained if they showed ≥60% pairwise sequence identity to any query and a full-length protein length ≥400 amino acids. The highest observed E-value among retained sequences was 7×10^-146^. A total of 46 sequences were obtained using these criteria. Multiple sequence alignment was generated with MUSCLE 5.3 (linux64) ^32^. A phylogenetic tree was inferred with IQ-TREE v3.0.1 ^33^ and visualized with iTOL v7 ^34^.

### Synthesis, recombinant expression, and purification of selected candidates

The open reading frames (ORF) encoding the selected urethanase candidates were codon-optimized for expression in *E. coli*, synthesized, and cloned into pETM11 by Genscript ^35^. The final construct contained an N-terminal 6xHis tag suitable for purification by immobilized metal affinity chromatography (IMAC) and a protease recognition site. Transformed BL21 (DE3) cells (Invitrogen) were grown overnight in 10 mL Lysogeny Broth (LB) media supplemented with kanamycin (50 µg mL^-1^). The LB medium (1 L) supplemented with kanamycin (50 µg mL^-1^) was inoculated with the *E. coli* preculture to reach a final optical density measured at 600 nm (OD600) of 0.025. Flasks were grown at 37 °C (200 rpm) to an optical density OD600 of 0.6–0.8 (Innova® 44, New Brunswick Scientific™). The culture was chilled at 4 °C for 30 min, followed by induction of protein expression with 0.1 mM isopropyl β-D-1-thiogalactopyranosid (IPTG). It was cultivated at 18 °C with shaking (200 rpm) for 16 h. Subsequently, cells were harvested by centrifugation (5,000 x g), and pellets were stored at -20 °C until further use.

Cell pellets from 1 L culture were resuspended in 3X volumes of lysis buffer (Buffer A: 10 mM Tris/HCl pH 8, 400 mM NaCl), containing 0.1 mg/mL DNase I (Sigma-Aldrich, Germany). Cells were lysed with Sonication (5s ON; 9s OFF; 1 minute total ON; 3 cycles). The cellular fraction was separated from the soluble lysate by centrifugation at 30,000 g, 4°C for 45 min (Sorvall LYNX 6000, Thermo Scientific). The cell-free extract (CFE) was filtered through 0.45 µm filters (Merck) and loaded onto a 5 mL Ni-NTA column (HistTrap, Cytiva), previously equilibrated with Buffer A, using an ÄKTA System (Cytiva). After complete sample loading, the column was washed with washing buffer (Buffer B: 10 mM Tris/HCl pH 8, 400 mM NaCl, 10 mM imidazole) until the absorbance at 280 nm returned to baseline levels. To elute bound proteins, a linear gradient of imidazole (10–300 mM) was applied to the column using elution buffer (Buffer C: 10 mM Tris/HCl pH 8, 400 mM NaCl, 300 mM imidazole). Fractions containing the protein of interest were analyzed by sodium dodecyl sulfate-polyacrylamide gel electrophoresis (SDS-PAGE) and then pooled and concentrated with Amicon® Ultra-15 centrifugal filters with a 30 kDa molecular weight cutoff (Utlracel®-30K; Merck-Millipore Ltd., Cork, Ireland). Buffer exchange was performed in storage buffer (Buffer D: 50 mM HEPES, 100 mM NaCl; pH 8) on a Desalting PD-10 column (Cytiva). Prior to crystallization, u15 was subjected to size-exclusion chromatography on a Superdex 200 16/60 (Cytiva) column and concentrated to 30 mg/mL using an Amicon® Ultra-15, then stored at -80 °C.

### Crystallization, Data collection, and Structure determination

The construct u15 was crystallized using sitting-drop vapor diffusion plates (Swissci, Molecular Dimensions) with a protein concentration at 31 mg/mL. Different commercial kits were used to screen initial hits by using a Mosquito Xtal3 (SPT Labtech) robot with different ratios between protein and reservoir (1:1; 1:2; 2:1). Multiple crystal hits were obtained, yielding microcrystals and smaller single crystals. The best crystallization condition was found in the BCS screen (Molecular Dimension) containing 0.2 M Ammonium sulfate, 0.05 M Magnesium sulfate heptahydrate, 0.1 M BICINE pH 9.0, and 20 % v/v PEG Smear Medium (PEG 3350, 4000, 2000, 5000 MME). Cryoprotection was achieved with 25% (v/v) ethylene glycol in mother liquor, prior to flash-freezing at 100 K in liquid N2. X-ray diffraction data were collected at the Deutsches Elektronen-Synchrotron, Hamburg, Germany (DESY). The data was processed at the beamline using EDNA with space group P 21 21 2 ^36^. The molecular replacement solution was found with Phaser using an AlphaFold prediction of SP3 as a search model ^37^. The crystal structure was refined and manually optimized using phenix.refine and Coot37 ^38^. Ramachandran plots and further validation were performed with Molprobity^39^. Molecular figures were produced using Pymol ^40^. Model and structure factors are deposited in the Protein Data Bank (PDB) with PDB ID 9SK0.

### Chromophore synthesis

The detailed procedure of Chromophore synthesis and characterization has been described elsewhere. ^41^

### DUE-MDA synthesis

The detailed procedure of DUE-MDA synthesis and characterization has been described elsewhere. ^27^

### Preparation of Flexible PUR Foam

The detailed procedure of Flexible PUR Foam preparation has been described elsewhere. ^29^ After foaming and curing, the material was processed through a meat grinder.

### Molecular Docking and Molecular Dynamics simulations

The crystal structure of the u15 enzyme was used as the framework for all simulations. We refined the structure by removing all ethylene glycol (EDO) molecules, chain B atoms, and crystallographic water molecules located more than 6 Å from any atom of chain A. Next, we set the protonation state of all residues at pH 9.0 using PROPKA 3.5.0 ^42^. The catalytic Lys78 was predicted to be neutral (-NH2), consistent with its buried position within the active site cavity. In this environment, Lys78 is positioned near two serine residues, Ser154 and Ser172 (≈ 2.9 Å apart), which can accept H-bonds from it, and close to cis-Ser153 (≈ 2.8 Å apart), which can donate an H-bond to Lys78. We conducted a visual inspection of all titratable amino acids and their surroundings to carefully check and validate the chosen protonation states. Among the sixteen existing histidine residues, four (His94, His202, His296, His419) were Nδ-protonated, while the remaining twelve were Nε-protonated. Hydrogen atoms were added to the structure using the XLEaP module of the AMBER 18 package ^43^.

We generated the u15:DUE-MDA complex by molecular docking using GOLD (Genetic Optimization Ligand Docking) ^44^. The DUE-MDA ligand was built and parameterized following the same procedure described in our previous work ^26^. Docking was performed with the binding region defined as a 30 Å radius sphere centered on the Oγ atom of the catalytic Ser177, with distance constraints (1.5-3.0 Å, spring constant of 5.0) between the carbonyl oxygen of the DUE-MDA’s urethane group and the NH backbone groups of Ile174 and Gly175. These constraints ensured proper placement of the carbamate moiety within the oxyanion hole and in close proximity to the nucleophilic Ser177. The poses were scored using the GoldScore fitness function ^44^, and convergence was assumed when the top three solutions differed by less than 1.5 Å RMSD (Root Mean Square Deviation). The docking solutions mainly adopted two distinct poses that differed in the relative orientation of the target carbamate bond of DUE-MDA with respect to the catalytic Ser177. We thus selected the highest-scoring complex from each pose for subsequent molecular dynamics (MD) simulations. Each u15:DUE-MDA complex was neutralized with Na+ counterions and solvated in a rectangular box of TIP3P water molecules ^45^ within a radius of 12 Å from the surface of the complex, yielding approximately 60,000 atoms. All-atom simulations were carried out with the GROMACS software (version 2024.4) ^46^ using the AMBER ff14SB protein force field ^47^. Each solvated u15:DUE-MDA system was submitted to a multi-step energy minimization procedure using the steepest descent algorithm. The minimized systems were then submitted to MD simulations, which started with a heating phase (100 ps, NVT ensemble), where each system was gradually warmed to 40 °C using the V-rescale thermostat ^48^, while solute atoms were kept fixed. We then conducted a 2 ns NPT equilibration phase to adjust the density of the systems to 1.0 bar using the Berendsen barostat ^48^, with all solute atoms still restrained. Subsequently, we performed a 2 ns NPT equilibration phase to relax the overall structure of the enzyme, while fixing the position of the active-site residues (Lys78, cis-Ser153, Ile174, Gly175, and Ser177) and the atoms forming the substrate’s urethane bond. Finally, we conducted a 400 ns NPT production phase without any restraints. Bonds involving hydrogen atoms were constrained with the LINCS algorithm ^49^, using a 2 fs integration timestep, a Verlet cut-off scheme, and a non-bonded cut-off value of 10 Å. Periodic boundary conditions were considered in all phases, and the non-bonded Coulombic interactions were treated using the Particle Mesh Ewald scheme ^50^. Coordinates were saved every 100 ps. Each complex was simulated in triplicate with distinct initial velocities, resulting in a total production time of 1.2 μs per system (Pose A and Pose B).

We used an identical procedure to generate and simulate the u15:MUE-MDA complex. Thus, we constructed the MUE-MDA ligand, parameterized it, docked it to the u15’s active site, and simulated the resulting complex for 400 ns in triplicate under identical conditions.

The trajectories of each set of triplicates were concatenated into a single trajectory for analysis. RMSD and root-mean-square fluctuation (RMSF) values were computed using GROMACS’s gmx rms and gmx rmsf tools, respectively. Interatomic distances were analyzed using the gmx distance as a function of simulation time. We used Visual Molecular Dynamics (VMD) ^51^ and Pymol software to visualize the trajectories and conduct structural analysis. ^40^

### Biochemical characterization

#### Determination of pH optima

Enzymatic activity was determined by an absorbance-based method using the Chromophore (Figure S3) with a cleavable urethane. The assay mixture consisted of 2 mM chromophore, 0.05 µM protein, and a series of buffers with different pH (pH 4-6 citrate buffer; pH 7-8 Tris HCl buffer; pH 9 Bicine buffer, and pH 10 CHES buffer). Measurements were carried out at 40 °C, with a 96-well plate (Greiner), using a Synergy HT plate reader (BioTek Inc.), and product formation was followed by monitoring the absorbance at 435 nm from 1 to 5 minutes. Each point was assayed in technical duplicate. The extinction coefficient of the reaction product has been determined by building a standard curve from 6-800 µM and measuring absorbance at 435 nm, normalized with path length.

#### Thermal Unfolding analysis

Unfolding profiles were measured on Prometeus Panta (NanoTemper). Purified protein (0.1 mg/ml final concentration) was assayed in different conditions. The temperature was increased 1.5 °C per minute from 20 °C to 90 °C. Apparent protein melting temperature (Tm app) was estimated on the Panta analysis software from NanoTemper using the initial ratio 350nm/330nm. .

#### Steady-state kinetics on Chromophore

The assay mixture consisted of 0-2 mM chromophore, 0.05 µM enzyme in 50 mM Bicine pH 9. Measurements were carried out at 40 °C, with a 96-well plate (Greiner), using a Synergy HT plate reader (BioTek Inc.), and product formation was followed by monitoring the absorbance at 435 nm from 1 to 5 minutes. Each point was assayed in technical triplicate. The extinction coefficient of the reaction product has been determined by building a standard curve from 6-800 µM and measuring absorbance at 435 nm, normalized with path length. Data treatment and statistics were done with MS Excel and GraphPad Prism 10.

#### Steady-state kinetics on DUE-MDA

Reaction mixtures (200 µL), containing 10-125 nM enzyme and 0.031–1.2 mM DUE-MDA (stock in pure Ethanol, resulting in a maximum 5% (v/v) solvent in the reaction mixture) in 50 mM Sodium Phosphate buffer (pH 8), were carried out in a benchtop thermomixer at 40 °C with shaking (800 rpm). In each case, the condition S0 >> E0 was fulfilled. After 5-10 min, the samples were quenched with a 10x dilution in pure ACN and filtered (0.45µm) for HPLC analysis. The presence of 90% pure ACN is sufficient to inactivate the enzymes by unfolding. Chromatographic separation was carried out using Accucore™ C18 (150 mm length x 4.6 mm diameter) HPLC Column (Thermo Fisher Scientific). The mobile phase consisted of 100% acetonitrile (ACN, Solvent A) and MilliQ Water (Solvent B). The flow rate was set to 2.2 mL/min, and the elution was performed under isocratic conditions at 40%/60% (SolventA/SolventB) for 6.5 minutes. The injected volume was 5 µL. Quantitation with UV-absorbance (240 nm) was performed by comparing the peak area of the compound of interest, DUE-MDA, MUE-MDA and 4,4-MDA,to the respective compound standard curve. Each point was assayed in technical duplicate. Data treatment and quantification were done with Chromeleon, MS Excel, and GraphPad Prism 10.

#### Progress curve on DUE-MDA

Degradation of DUE-MDA over time was measured in a reaction mixture (200µL) of 0.1 µM enzyme and 1 mM DUE-MDA (stock in 96% Ethanol, resulting in a maximum 5% (v/v) solvent in the reaction mixture), 50 mM Sodium Phosphate buffer (pH 8), in a benchtop thermomixer at 40 °C with shaking (800 rpm). Reactions were quenched by a 10x dilution in pure ACN after 15, 30, 45, 60, 90, and 120 minutes. Samples were filtered (0.45 µm) and analyzed by RP- HPLC as described above. The injected volume was 5 µL. Quantification and data treatment was performed as described above.

#### Progress curve on DUE-TDA

Degradation of DUE-TDA over time was measured in a reaction mixture (1 mL) of 0.1 µM enzyme and 1 mM DUE-TDA (stock in pure Ethanol, resulting in a maximum 5% (v/v) solvent in the reaction mixture), 50 mM Sodium Phosphate buffer (pH 8), in a benchtop thermomixer at 40 °C with shaking (800 rpm). Reactions were quenched by a 1:1 dilution in pure ACN after 30, 60, 90, and 120 minutes. Samples were filtered (0.45 µm) and analyzed by RP- HPLC. Chromatographic separation was carried out using Accucore™ C18 (150 mm length x 4.6 mm diameter) HPLC Column (Thermo Fisher Scientific). The mobile phase consisted of ACN (Solvent A) and Milli-Q water (Solvent B). The flow rate was set to 2.2 mL/min. The elution was initiated under isocratic conditions at 20%/80% (Solvent A/Solvent B) for 2.5 minutes, followed by a gradient increase of ACN until 4 minutes where it reached to 95%/5% (Solvent A/Solvent B) over 1 minute. Then a 0.5 minute gradient return to 20%/80% and a final hold for 1 minute. The total run time is 6.5 minutes. The injected volume was 5 µL. Quantitation with UV-absorbance (240 nm) was performed by comparing the peak area of the compound of interest, 2,4-TDA, to the respective compound standard curve. Each point was assayed in technical duplicate. Data treatment was performed as described above.

#### Endpoint measurement of hydrolytic activity on Flexible PUR Foam

The reaction mixture (1 mL) contained 25 mg/ml Flexible PUR Foam, 5 µM enzyme, 50 mM Sodium Phosphate buffer (pH 8). The reaction was carried out in a benchtop thermomixer at 40 °C with shaking (800 rpm) and was quenched after 24 and 48 hours by a 1:1 dilution in pure ACN. Samples were filtered (0.45 µm) and analyzed by RP- HPLC as described above. The injected volume was 20 µL. Quantitation with UV-absorbance (240 nm) was performed by comparing the peak area of the compound of interest, 2,4-TDA and 2,6-TDA, to the respective compound standard curve. Each point was assayed in technical triplicate. Data treatment was performed as described above.

### Results and Discussion Homology Search

New urethanase candidates were initially identified through an homology-guided search using UMG-SP1, UMG-SP2, and UMG-SP3 as queries ^24^. In the first round, we pooled sequences with up to ∼50% sequence identity (SI) to the queries and selected five putative urethanase sequences for expression and purification. As neither of these homologs showed both detectable activity and robust expression, we then narrowed the search space to sequences between 60% and 80% SI. We then constructed a phylogenetic tree of homologs sharing ≥60% identity with at least one query (Figure 1A) and selected four representatives (u9–u12) to maximize phylogenetic diversity within the cluster, but that shared less than 80% sequence identity each other. The present study does not include u11, which will be described elsewhere. Notably, u10 was strongly expressed and exhibited high thermostability, motivating a third, u10-based, round of homology search that yielded candidates u13–u17. The final panel displayed ∼50% identity to UMG-SP2/SP3 and ∼70% identity to UMG-SP1 (Figure 1B, Figure S2). Two pairs, u10/u13 and u15/u16, were especially closely related (approx. 90% SI). We included the sequences with high SI, as we experienced differences in expression levels and solubility. All selected sequences originate from Sphingomonas strains, a genus known for its ability to catabolize aromatic compounds.^52^ Among the set, u17 was the most representative, showing the highest identity to UMG-SP1 and to the remainder of the candidates. All candidate genes were codon optimized and cloned into the pETM11 vector for recombinant expression in *E.coli* BL21 DE3.

**Figure 1.**
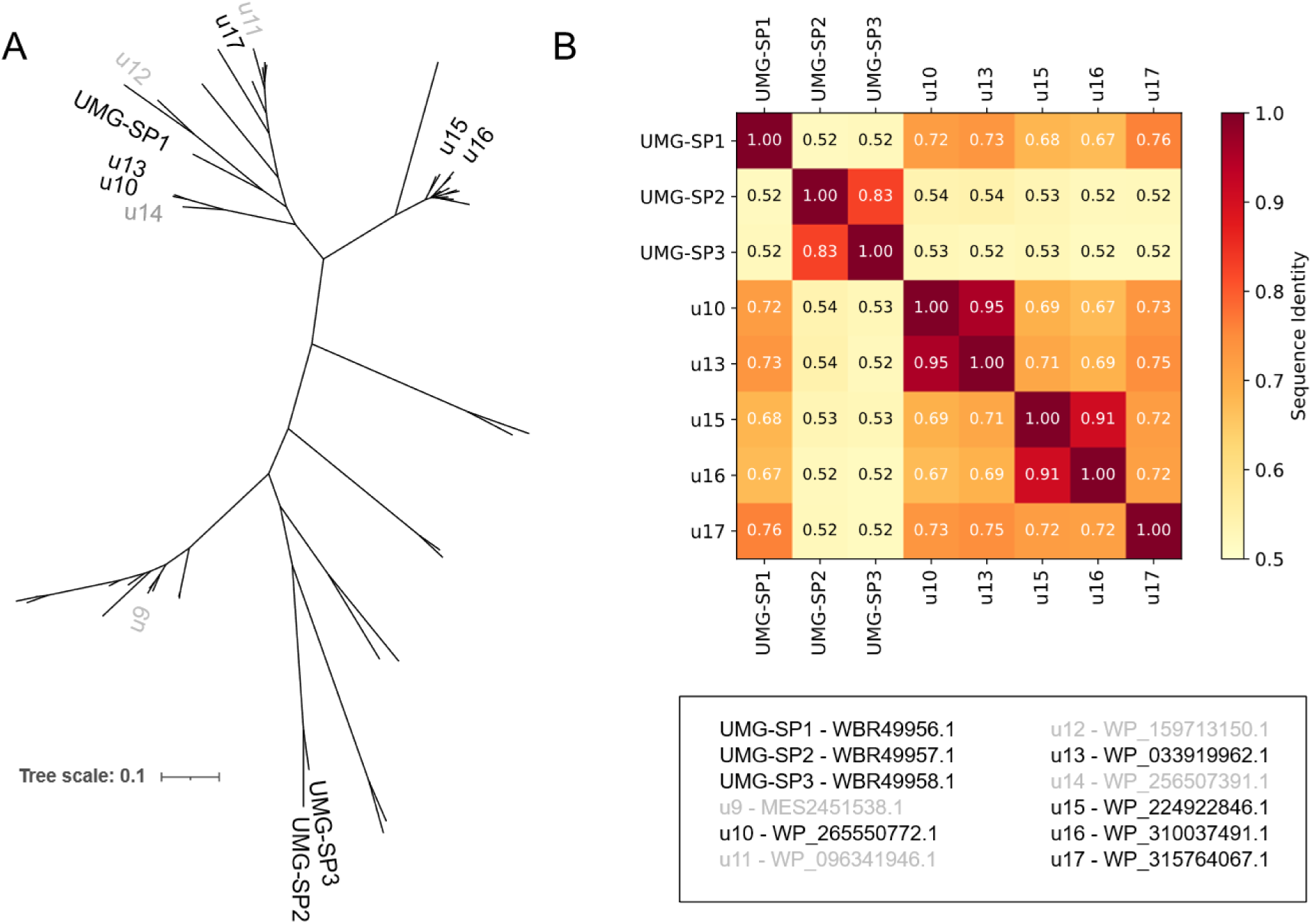
Phylogenetic and sequence-identity analysis of selected urethanase candidates. A) Phylogenetic tree constructed using u10-17 sequences together with known urethanases UMG-SP1-3 for a visual overview. Grey candidates either did not express in *E. coli* BL21(DE3) (u14) or failed to show higher thermostability than u10 (u9-u12). B) Sequence comparison based on sequence identity between the selected candidates u10-17 and the previously characterized UMG-SP1-3, the heat map describes the highest SI percentage as dark red (SI 100%) to the lowest yellow (SI 50%). The NCBI reference codes are as follows: u9 (MES2451538.1, *Sphingomonas* Bacterium Eh05), u10 (WP_265550772.1, *Sphingomonas* sp. S1-29), u11 (WP_096341946.1, *Sphingomonas spermidinifaciens*), u12 (WP_159713150.1, *Sphingomonas* sp. AX6), u13 (WP_033919962.1, *Sphingomonas* sp. 37zxx, u14 (WP_256507391.1, *Sphingomonas qomolangmaensis*), u15 (WP_224922846.1, *Sphingomonas* sp. R647), u16 (WP_310037491.1, *Sphingomonas* sp. BE123), u17 (WP_315764067.1, *Sphingomonas* sp. Y38-1Y), UMG-SP1 (WBR49956.1, Uncultured Bacterium), UMG-SP2 (WBR49957.1, Uncultured Bacterium), UMG-SP3 (WBR49958.1, Uncultured Bacterium).

### Biochemical characterization

The purified enzymes were initially screened on a synthesized chromophore to quantify carbamate hydrolytic activity by monitoring product release at 435 nm (Figure S3). This assay offers the simplicity and high-throughput of an absorbance-based screening method while using a sterically demanding substrate. Consequently, it enables the selection of enzymes that not only possess carbamate cleaving activity but are also capable of accommodating bulky molecules within their active sites, thereby mimicking the steric constraints imposed by polyurethane fragments. All urethanases tested exhibited maximal catalytic activity at pH 9, except for u13, which maintained stable activity across the entire pH range of 7 to 10 (Figure 2A). This trend is consistent with the known behavior of signature amidases, where alkaline conditions favor the deprotonation state of key active site residues, enhancing nucleophilic attack on the carbamate carbonyl and facilitating catalysis ^53,54^. This behavior was also observed in SP1 ^24,28^, which showed optimal activity at pH 8–9, thereby validating both the experimental setup and the observed alkaline activity profiles of the newly identified enzymes. Thermal stability was assessed using nano differential scanning fluorimetry (NanoDSF) under varying pH values and acetonitrile (ACN) concentrations (Figure 2B-C). All enzymes maintained stability in the pH range 7 to 10, with u17 and u16 representing the most and least thermostable variants (Tm app = 67 °C and 50 °C, respectively) (Figure 2B). Regarding solvent tolerance, comparable behavior was observed upon exposure to ACN at pH 9, where unfolding tendency increased with ACN concentration (1–20% v/v). At 20% ACN, all urethanases demonstrated substantial thermal loss between (Figure 2C). Under this condition, complete unfolding precluded thermal profile determination for u10. In both thermal stability assays, SP1 served as the reference enzyme. Defining optimal reaction parameters was a prerequisite for determining the kinetic constants for turnover under steady-state conditions using the Chromophore. All urethanases demonstrated notable activity, with turnovers ranging from 2.6 s⁻¹ to 15 s⁻¹ (Figure 2D, Table 1). The enzymes exhibited modest substrate affinity, with Michaelis–Menten constants (K_M_) in the millimolar range. As a result, specificity constants span from 2 (u17) to 12 s^-1^ mM^-1^ (u10) (Table 1). Importantly, the highest k_cat_/K_M_ values were observed for u10 and for the control enzyme SP1, highlighting their comparatively high catalytic efficiency under the tested conditions and reinforcing SP1 as a reliable benchmark for functional comparison. High K_M_ values are likely attributable to the Chromophore, being a nonnatural substrate with substantial steric bulk, which may hinder optimal accommodation within the catalytic pocket and thus hamper saturation under assay conditions. Moreover, for u15 and u16 substrate saturation was not fully reached within the tested concentration range and consequently the fitted K_M_ exceeded the highest substrate concentration assayed. As a result, this values should be interpreted as an estimate.

**Figure 2.**
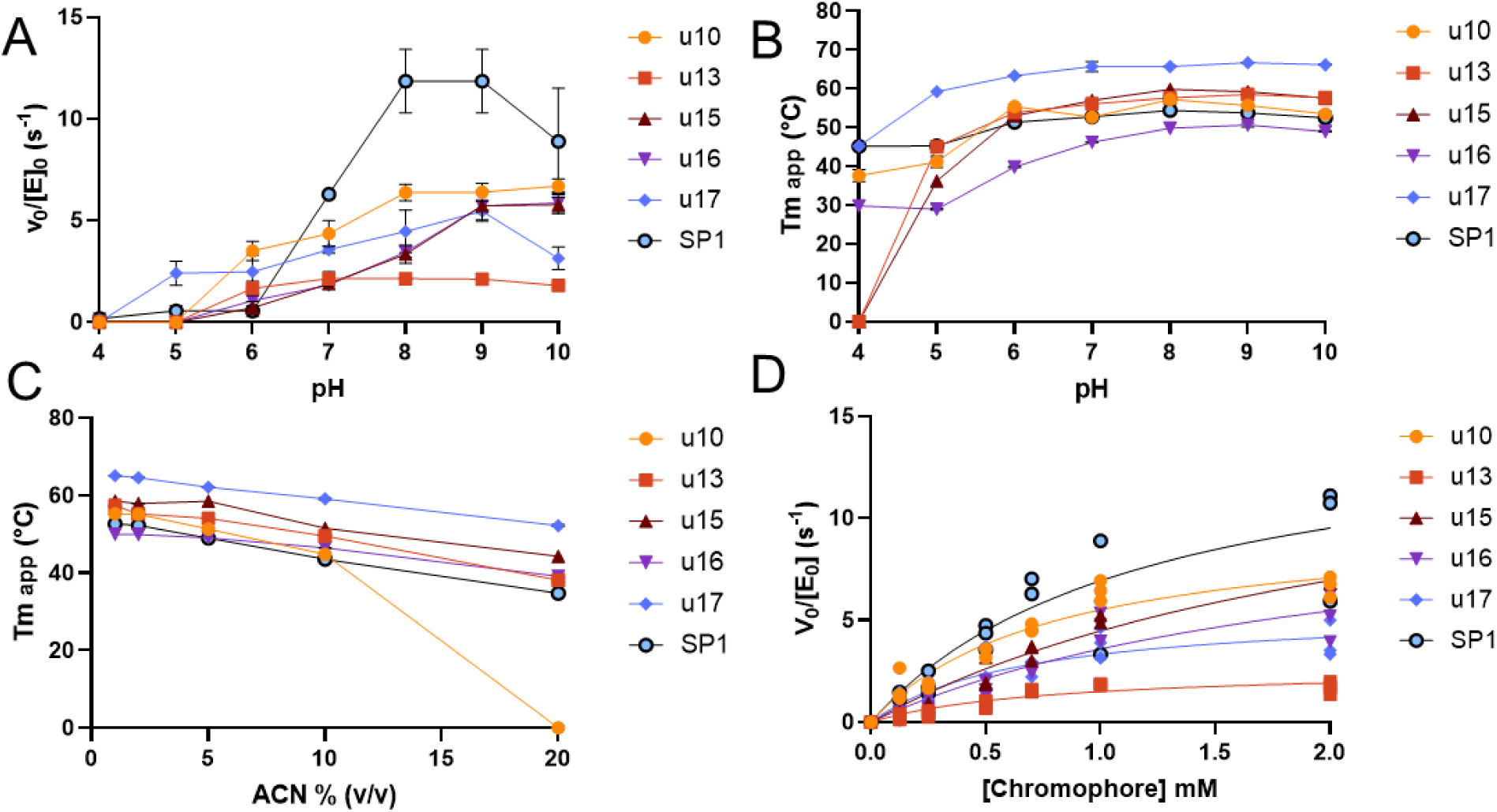
Biochemical characterization of u10,u13, u15-17 and SP1. A) Specific activity as a function of pH of u10, u13, u15, u16, u17, and SP1. B) Apparent unfolding temperature of urethanases assayed in the pH range between 4 and 10. C) Apparent unfolding temperature (Tm app) of the tested candidates under different acetonitrile concentrations (1-20% v/v). D) Michaelis-Menten curves obtained on the Chromophore. Data points represent the Mean and standard deviation from three independent measurements (n=3) .

**Table 1.**
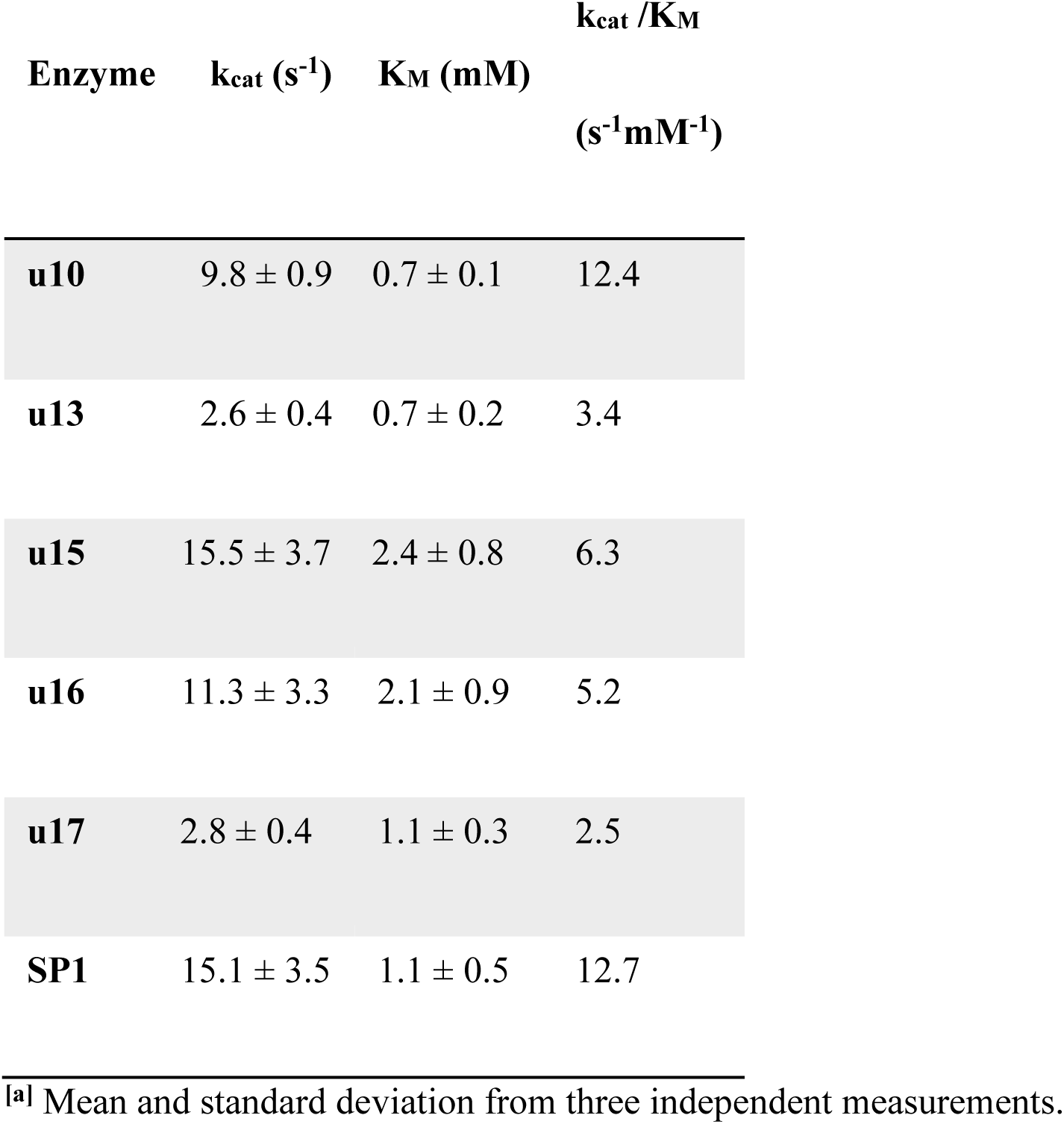
Steady-state kinetic parameters on Chromophore^[a]^.

### Hydrolysis of a Polyurethane Model Substrate

To evaluate urethanase activity towards polyurethane (PUR)-like structures, we designed and synthesized a soluble model substrate for polyether-based PUR, DUE-MDA (Figure S1A) ^27^. This compound was accepted as a substrate by all enzymes investigated, and the reaction progress was monitored by HPLC (Figure S4). The predominant reaction was the cleavage of a single urethane bond, resulting in the release of a diethylene glycol methyl ether substituent and the formation of a mono-substituted MDA-urethane derivative, hereafter referred to as MUE-MDA ^27^.

Steady-state kinetic analysis revealed that turnover numbers (k_cat_) for DUE-MDA ranged from below 1 s^−1^ for variant u10 to 14.6 s^−1^ for variant u15 (Figure 3; Table 2). Michaelis–Menten constants (K_M_) were generally similar among the tested enzymes, with the exception of u10 that did not reach saturation under the tested conditions and was therefore excluded from direct comparison of individual catalytic performances. For most enzymes analyzed, catalytic efficiency (k_cat_/K_M_) reflected a relatively high turnover and modest substrate affinity, not far from what was expected ^55,56^. Candidates like u15, u16, and u17 emerged as the most efficient catalysts, outperforming u10 and u13. In parallel, kinetic benchmarks were established by comparison with the closest reference enzyme, SP1, which displayed a turnover number of approximately 4.5 s^−1^ and a k_cat_/K_M_ around 15 s^−1^ mM^−1^ (Table 2).

**Figure 3.**
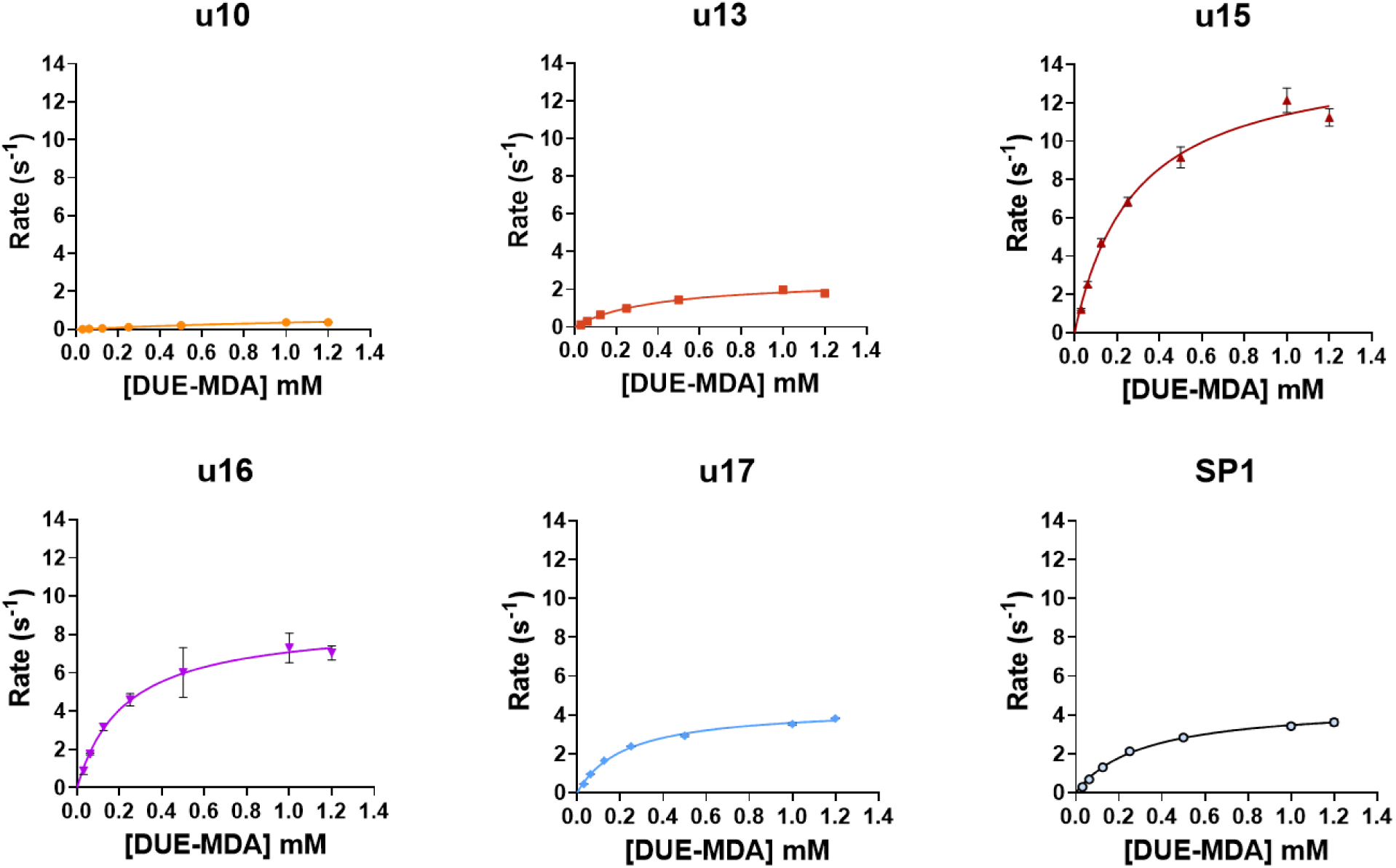
Steady-state kinetics on DUE-MDA. Michaelis-Menten curves of u10, u13, u15-u17, and SP1 obtained by monitoring DUE-MDA hydrolysis. Each data point represents the mean and standard deviation from three independent measurements (n=3).

**Table 2.** Steady-state kinetic parameters on DUE-MDA.^[a]^.

| **Enzyme** | **$k_{\text{cat}}$ (s<sup>-1</sup>)** | **$K_{\text{M}}$ (mM)** | **$k_{\text{cat}}/K_{\text{M}}$**<br>**(s<sup>-1</sup>mM<sup>-1</sup>)** |
| --- | --- | --- | --- |
| **u10** | n/a | n/a | n/a |
| **u13** | 2.5 ± 0.2 | 0.4 ± 0.08 | 6.5 |
| **u15** | 14.6 ± 0.7 | 0.3 ± 0.04 | 51.8 |
| **u16** | 8.7 ± 0.2 | 0.2 ± 0.02 | 38.3 |
| **u17** | 4.4 ± 0.1 | 0.2 ± 0.02 | 19.7 |
| **SP1** | 4.5 ± 0.1 | 0.3 ± 0.03 | 14.8 |

<sup>[a]</sup> Mean and standard deviation from three independent measurements.

To complement steady-state characterization, time-course experiments were conducted to capture product profiles over two hours of reaction. These analyses confirmed the progressive depletion of DUE-MDA (Figure 4A) and the concomitant accumulation of MUE-MDA (Figure 4B, Figure S1A) and 4,4-MDA (Figure 4C, Figure S1A). After 120 minutes, concentrations of MUE-MDA (Figure 4B) reached 40 and 340 µM for u10 and u13, respectively, while higher levels were observed for the most active variants, reaching 1.3 mM (u15), 0.7 mM (u16), and 0.98 mM (u17). Notably, the formation of the fully hydrolyzed product 4,4-methylenedianiline, 4,4-MDA (Figure 4C), was strongly candidate-dependent: u17 and u15 were the only enzymes able to produce appreciable amounts, accumulating to 264 µM and 115 µM after 2 h, while in all other variants 4,4-MDA release remained confined to sub micromolar concentrations. Within the first 45 minutes, u15 produced higher amounts of both products, whereas u17 continued to generate products beyond this time point, consistent with its greater thermal stability. Together, these observations indicate that enzyme performance in the short assay is strongly influenced by differences in early turnover and stability, and they support u15 and u17 as promising candidates for further optimization.

**Figure 4.**
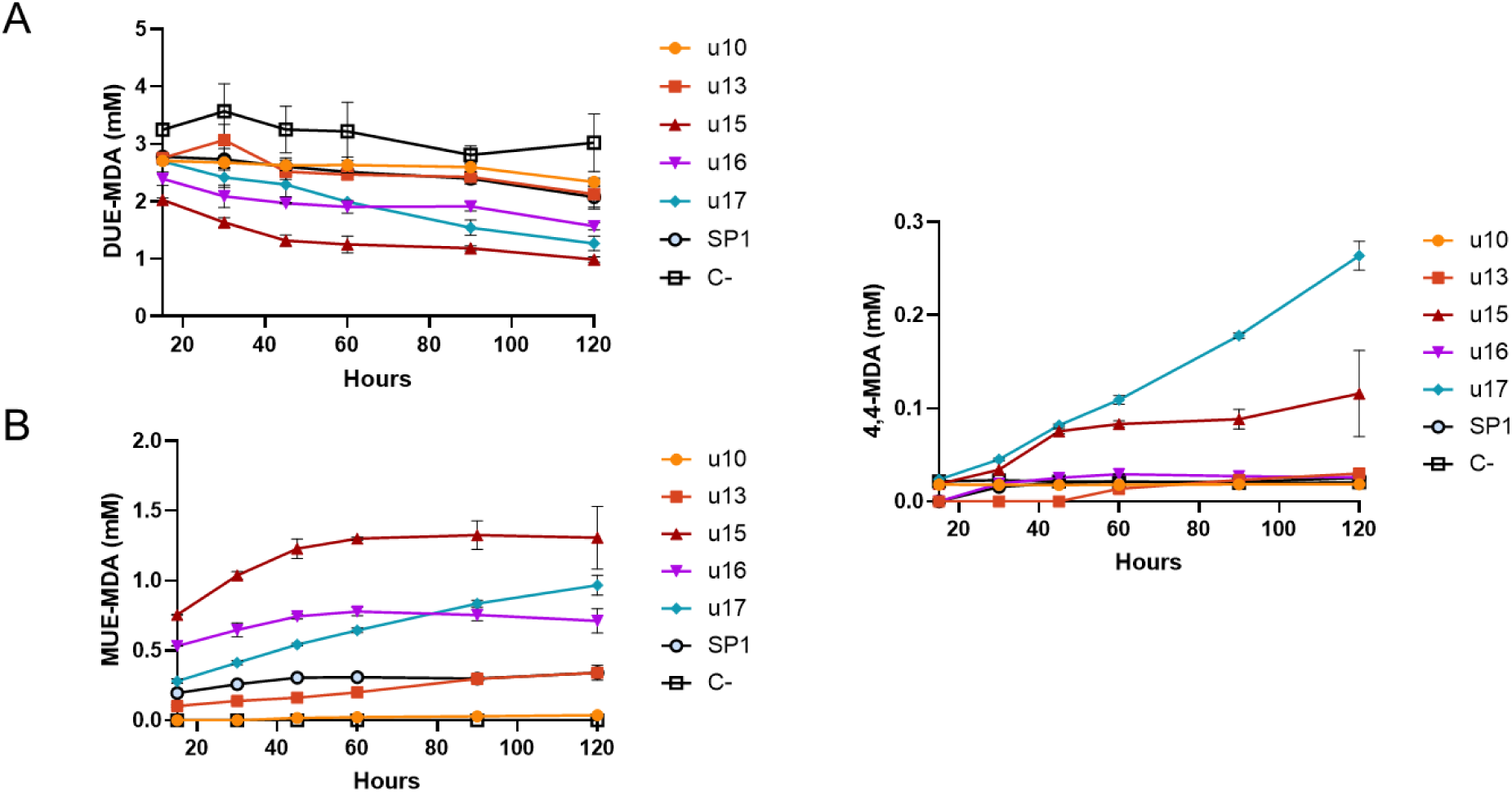
Progress curves of u10-17 and SP1. A), DUE-MDA hydrolysis monitored over the course of 2 hours. B) Release of mono-hydrolyzed MUE-MDA. C) Release of the fully hydrolyzed 4,4-MDA. The reaction consists of 0.1µM enzyme and 1mM DUE-MDA (maximum 5% Ethanol (v/v)) in 50 mM Sodium Phosphate buffer (pH 8), at 40 °C with 800 rpm shaking.

### Crystal Structure of u15

The crystal structure of u15 was determined at 2.05 Å resolution and determined in complex with ethylene glycol. Complete crystallographic statistics are provided in the Supporting Information (Table S1). The enzyme adopts a compact α/β-fold characteristic of the amidase signature (AS) family, consisting of twelve α-helices (αA–αL) arranged around a central β-sheet of ten β-strands (β1–β10) (Figure 5A). In line with its classification, u15 harbors the canonical AS catalytic triad composed of Ser177, cis-Ser153, and Lys78, embedded within the highly conserved amidase signature sequence (Figure 5B). The active site architecture is consistent with the reaction mechanism reported for related enzymes such as UMG-SP2 ^57^. During catalysis, Ser177 acts as the principal nucleophile, initiating attack on the substrate and forming a tetrahedral intermediate stabilized within an oxyanion hole generated by the backbone amide groups of, in the case of u15, Ile174 and Gly175. Proton transfer during the reaction cycle occurs between cis-Ser153 and Lys78, facilitating breakdown of the intermediate and completion of the hydrolytic process. Moreover, the catalytic triad is stabilized by Ser154 and Ser172 acting as bridging residues. The structure of u15 was determined in complex with ethylene glycol, a product of the DUE-MDA breakdown, and also used as a cryoprotectant prior to cryogenic freezing; the ligand is positioned within the active site, coordinated at a distance of 3.3 Å from the catalytic oxygen of Ser177, and is fully enclosed within the enzyme’s active site pocket (Figure 5B). The substrate binding pocket is delineated by α-helices I (residues 292-307) and K (residues 380-391) and loops 1–3 (L1 residues 74–101; L2 117–151; L3 187–212 and L4 352–384). The pocket is predominantly hydrophobic in character, creating a defined pathway that directs substrates toward the catalytic center (Figure 5C). Key residues, Leu303, Phe307, Thr300, Trp130, Leu383, Gln365, Asp129, and His296 protrude to define the structural contour of the binding area (Figure 5D).

**Figure 5.**
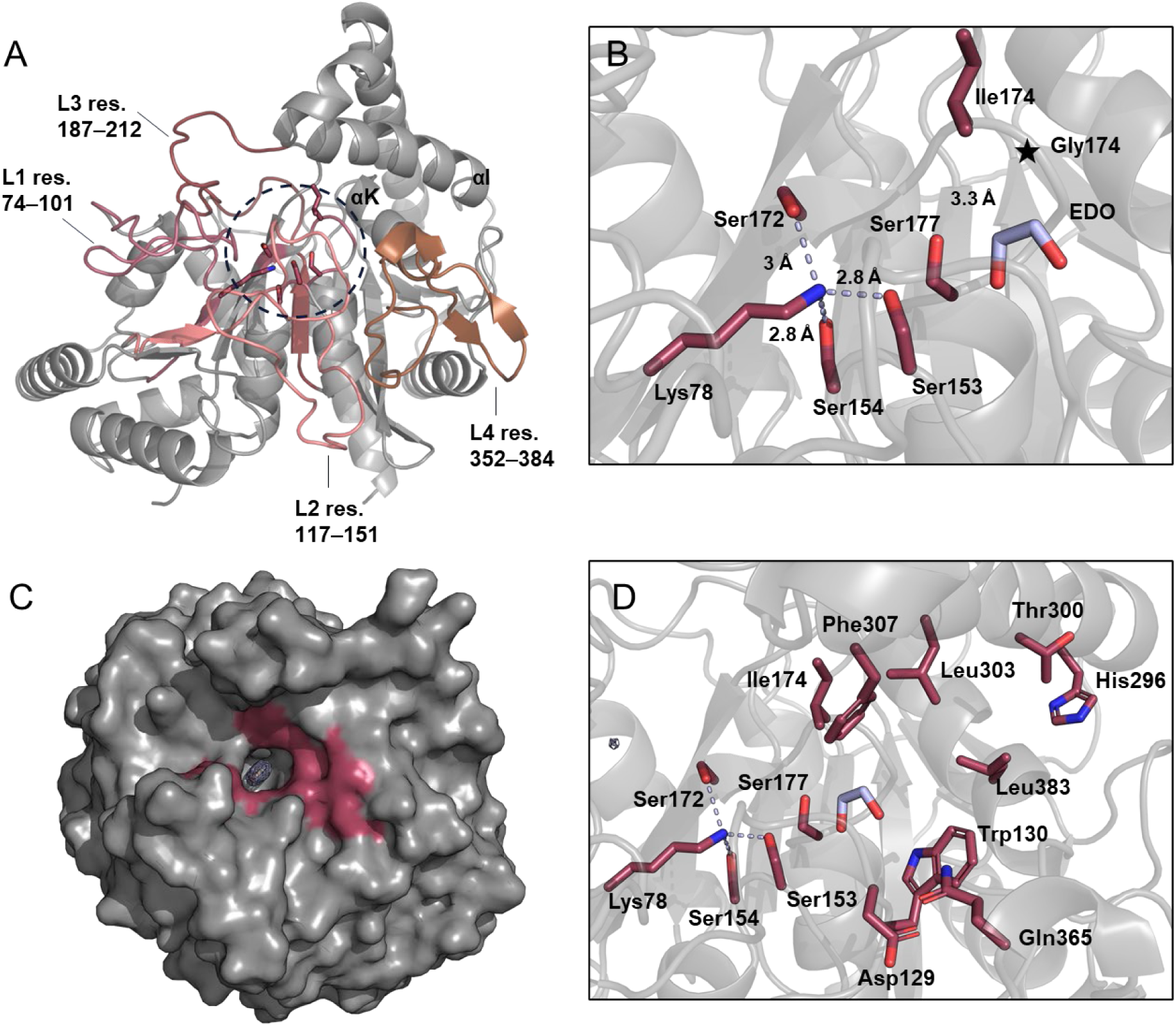
Crystal structure of urethanase-15 (PDB ID 9SK0). A) Overall structure of u15 showing the four loops and αI (residues 292-307) and αK (residues 380-391) surrounding the catalytic triad (circled with dashed line). B) Close-up of the catalytic triad of u15 Ser177, cisSer153 and Lys78. The oxyanion hole Ile172 and Gly174 (the latter shown as a star of the backbone) and the bridging residues Ser172 and Ser154. The structure is in complex with ethylene glycol (EDO), shown in lilac. C) u15 structure shown as surface representation in complex with ethylene glycol (lightblue). The active site and the substrate pocket are colored in raspberry. D) Close-up of the substrate binding pocket of u15. Dashed line represents Hydrogen bonds.

Structural homology searches using the Dali server revealed the closest matches to be urethanase UMG-SP3 (PDB 9FVF; RMSD 1.1 Å covering 429 aligned Cα atoms, Z score of 63.5, 55% sequence identity)^27^ and ClbL amidase from the colibactin gene cluster (PDB 8ES6; RMSD 1.7 Å over 407 aligned Cα atoms, Z score of 53.4, 36% sequence identity)^58^. UMG-SP1 (PDB 9GIZ; RMSD 1.2 Å over 374 aligned Cα atoms, Z-score 58.6, 67% sequence identity)^28^ was also identified as a close homolog, although several disordered regions in proximity to the active site limited its suitability for detailed structural comparison. Both UMG-SP3 and ClbL structures exhibit an overall architecture that is equivalent to u15, including a highly conserved hydrogen-bonding network around the catalytic triad. As in other AS family members, the substrate-binding pocket of u15 is predominantly hydrophobic and solvent-accessible ^28,57–59^. A comparative analysis with SP3 in complex with 4,4-MDA (PDB ID 9FVF), the product of complete DUE-MDA hydrolysis, was carried out to rationalize observed functional differences (Figure 6). While the core fold and active site geometry are preserved, notable local variations were identified in the substrate-binding pocket and in the conformation of L3 and L4. In u15, Phe307 replaces Leu308 in SP3 and Gln365 substitutes for SP3 Trp367 (Figure 6A). Analogous to the flexible behavior observed for L3 in SP3, where Arg209 undergoes substantial conformational rearrangement in response to ligand binding ^27^, u15 features Glu211 at the equivalent position, oriented outward the catalytic center and likely engaging in an induced fit mechanism upon ethylene glycol association (Figure 6B). A six-residue stretch in L4, spanning Pro362–Glu367 in u15 and Glu364–Lys369 in SP3, distinguishes this region structurally between the two proteins and underlies significant divergence in their local architecture (Figure 6C). These substitutions, particularly in the binding pocket and loops proximal to the catalytic site, likely influence substrate recognition and product release, thereby contributing to the distinct catalytic profiles observed for u15.

**Figure 6.**
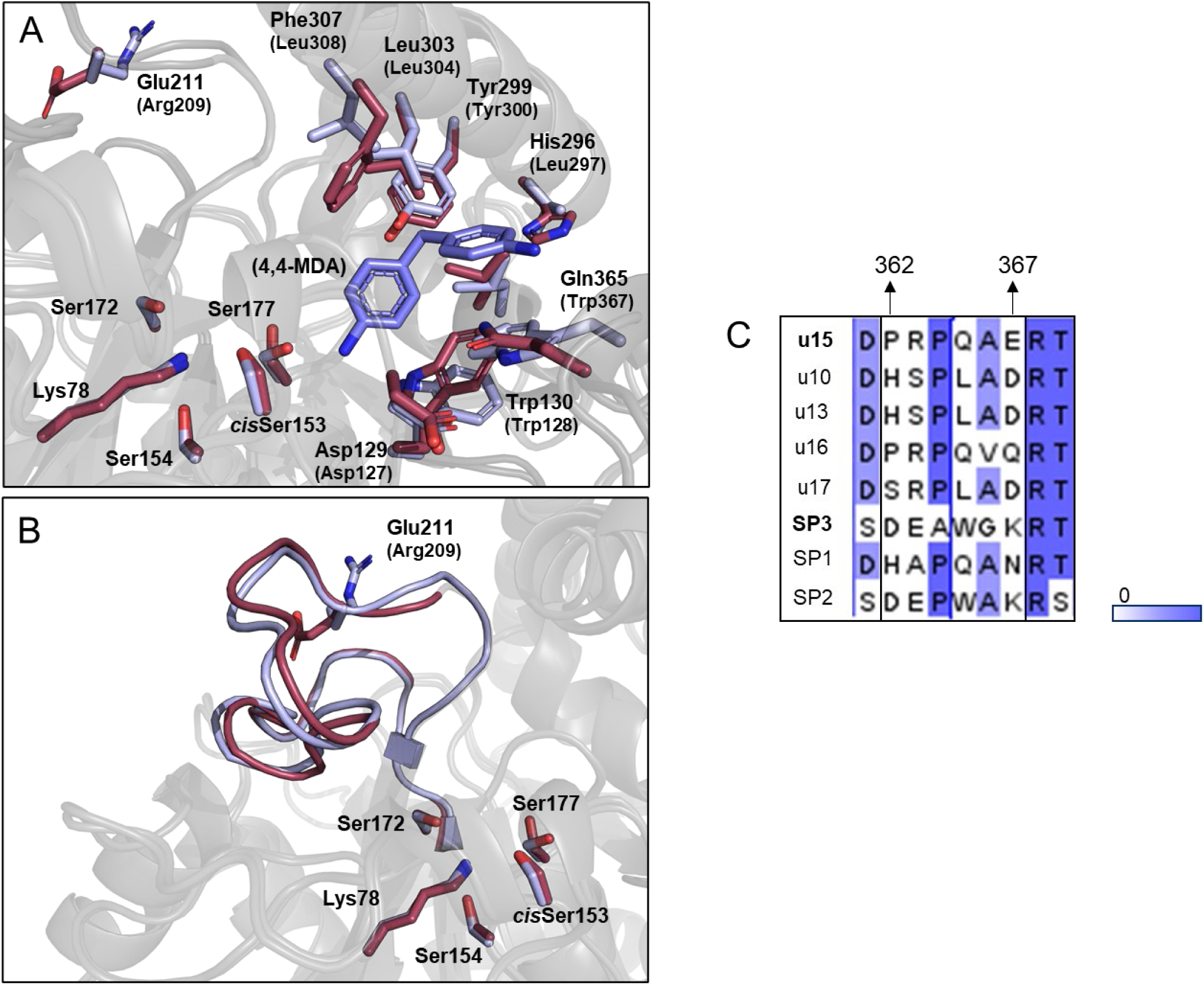
Structural comparison between u15 and SP3. A) Differences in the active site and binding pocket of u15 (PDB ID 9SK0 – colored bordeaux) and SP3 (PDB ID 9FVF – colored lightblue). Residues in brackets belong to SP3. B) Arrangement of Loop 3. res. 187–212 in u15 and res 189–214 in SP3. C) Comparison of Loop 4, with particular focus on a six-residue long stretch Pro362–Glu367 in u15. The sequences are colored by percentage of identity, with 0 being 0 % identical (white) and 1 being 100% identical (Blue).

To rationalize the catalytic differences observed between u15 and u17, we performed a comparative sequence and structural analysis (Figure S5). Extensive crystallization trials were conducted to resolve the crystal structure of u17. However, all obtained crystals exhibited a disordered active site, preventing reliable structural interpretation. To overcome this limitation and enable comparative analysis, an AlphaFold model of u17 was generated. U15 and u17 share 72% sequence identity, and structural alignment using Dali yielded an RMSD of 1 Å across 435 aligned Cα atoms (Z-score 68.9) (Figure S5A). The overall fold and topology of the two enzymes are highly similar, with only minor variations detected in the substrate-binding pocket. Most active site residues are conserved, except for two substitutions: Thr300 in u15 corresponds to Gly297 in u17, and Glu365 in u15 corresponds to Leu362 in u17. Overall, the binding pocket exhibits minimal variation, insufficient to explain the higher catalytic activity of u15 relative to u17 (Figure S5B). These findings demonstrate that differences in the binding pocket, together with mobile and flexible elements flanking the active site, can directly contribute to the broad substrate range characteristic of amidase-signature enzymes, motivating the need for future engineering campaigns. Yet, the structural comparison between u15 and u17, which share high sequence identity and nearly identical topology, suggests that structural similarity alone cannot account for the observed differences in catalytic efficiency. Taken together, these observations indicate that substrate adaptability in this enzyme family likely arises from a combination of active site architecture and dynamic properties such as local flexibility and stability under reaction conditions, two parameters that represent promising targets for rational optimization and for developing synergistic enzyme systems.

### Computational studies

Computational simulations are crucial for defining and rationalizing the intermolecular interactions that underlie catalytic activity. By providing detailed insight into these fundamental interactions, they enable the rational optimization of substrate affinity and reaction rate in future studies. To this end, we docked a DUE-MDA molecule into the crystal structure of u15, obtaining ten solutions that clustered into two distinct conformations differing in the relative orientation of the DUE-MDA’s carbamate bond relative to the catalytic Ser177. These were designated as Pose A and Pose B (Figure S6A). Pose A resembled, which was adopted by DUE-MDA in the UMG-SP2:DUE-MDA complex used previously to study its catalytic mechanism ^26^, where the 4,4-MDA moiety lies on the right side of Ser177’s side chain (Figure S6B). In this orientation, the 4,4-MDA moiety occupies the same region as in the crystal structure of the UMG-SP3:4,4-MDA (PDB ID: 9FZ1, Figure S6C) ^27^. This orientation is compatible with the cleavage of the carbamate ester bond, consistent with the mechanism previously described for the UMG-SP2-mediated hydrolysis of DUE-MDA. Conversely, in Pose B, the 4,4-MDA moiety is positioned on the left side of Ser177’s side chain (Figure S6D), an orientation we hypothesize may favor hydrolysis by the cleavage of the urethane C-N bond. Both poses may therefore coexist in the real system, each ultimately leading to urethane bond hydrolysis, although their relative stability is uncertain. For this reason, we selected the top-scoring complex from each pose for subsequent MD simulations to evaluate their relative stability and catalytic competence. Pose A proved more stable, with DUE-MDA exhibiting a smaller average RMSD (4.3 ± 0.9 Å vs. 5.5 ± 1.1 Å for Pose B). This stability was reflected in consistently shorter distances between the carbamate carbonyl oxygen and the backbone amides of Ile174 and Gly175 (≈ 2.1 Å in Pose A vs. 3.9-5.1 Å in Pose B) (Figure S7 and Table S2), indicating that the substrate remained lodged in the oxyanion hole cavity. Catalysis also appeared more favorable when DUE-MDA adopts Pose A, as the distance associated with the nucleophilic attack (i.e., between the Oγ of Ser177 and the carbamate carbonyl carbon) was more stable and shorter (3.3 ± 0.2 Å vs. 4.7 ± 1.5 Å, Figure S7 and Table S2). These findings suggest that u15 hydrolyses the DUE-MDA urethane bond through the same catalytic mechanism proposed for UMG-SP2 ^26^, involving release of an alcohol-leaving group followed by formation of a carbamic acid intermediate that spontaneously decomposes into an amine and CO₂. Based on these results, we conducted further analysis of substrate binding interactions using Pose A.

The MD simulations provided important insight into the role of key residues, corroborating hypotheses proposed in the previous section. As expected, Ser154 and Ser172 were crucial for stabilizing the catalytic triad, each consistently accepting a H-bond from Lys78 (≈ 2.1-2.2 Å, Figure 7A). This interaction enabled Lys78 to act as an H-bond acceptor from cis-Ser153 (1.8 ± 0.1 Å), which in turn accepted a H-bond from the nucleophilic Ser177 (2.9 ± 0.6 Å). Additionally, Ser154 further stabilized the catalytic triad by donating an H-bond to the backbone carbonyl oxygen of the nucleophilic Ser177 (2.0 ± 0.2 Å).

**Figure 7.**
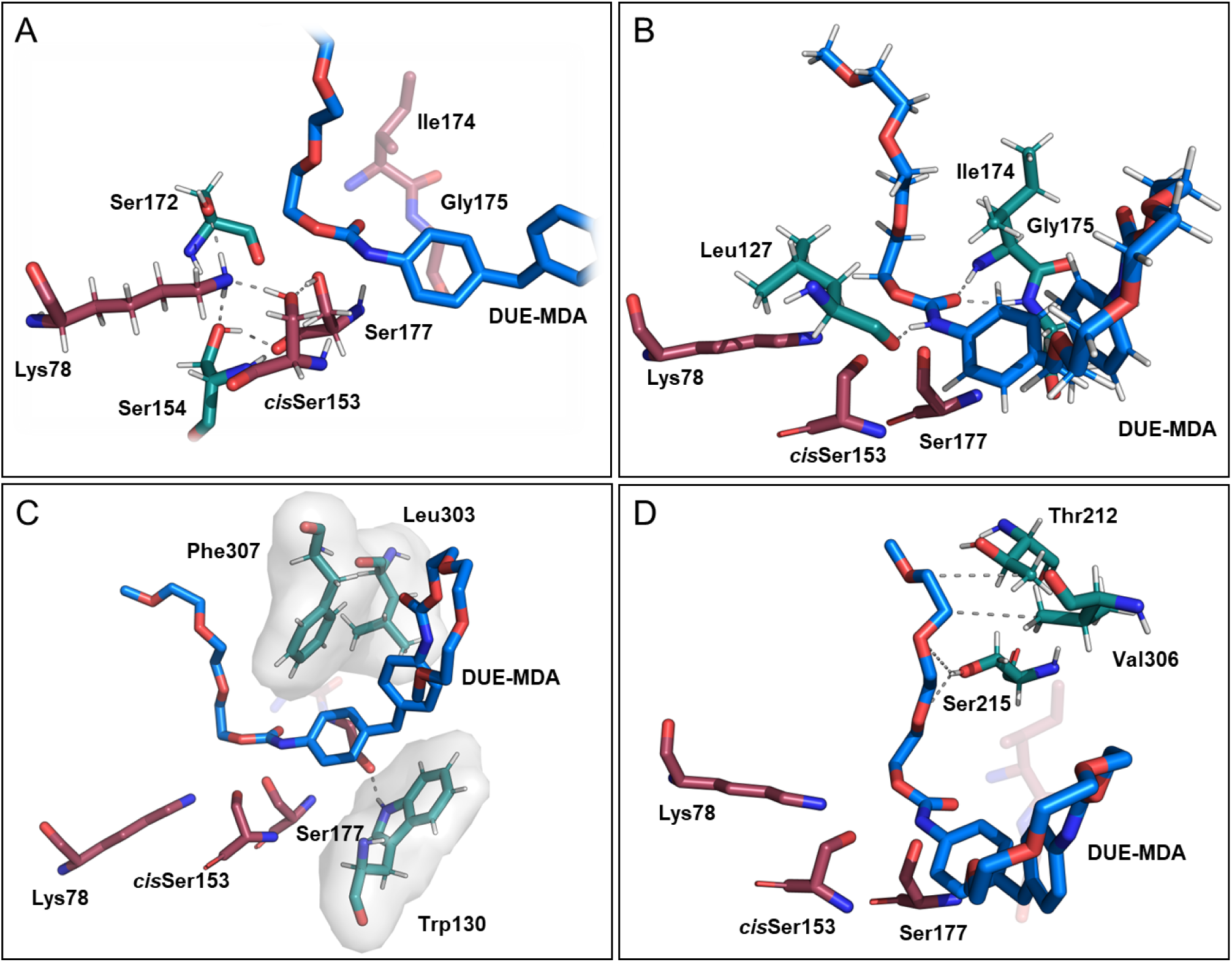
Molecular Dinamic simulation on u15:DUE-MDA complex. A) Ser154 and Ser172 (shown in green sticks) form persistent H-bonds with Lys78, contributing to the stability of the catalytic triad. B) Along with the oxyanion hole residues Ile174 and Gly175, Leu127 (shown in green sticks) maintains the DUE-MDA carbamate group properly oriented for catalysis. C) The hydrophobic triad formed by Trp130, Leu303, and Phe307 (shown as green sticks and grey surface) anchors the 4,4-MDA moiety of DUE-MDA. D) Thr212, Ser215, and Val306 transiently stabilize the substrate’s polyol chain attached to the target urethane bond. Hydrogen atoms are omitted in selected regions for clarity.

Along with the oxyanion hole amides, we identified Leu127 as a key residue in maintaining the proper orientation of the target carbamate group for nucleophilic attack, forming a persistent H-bond with the carbamate’s NH group (2.2 ± 0.4 Å, Figure 7B). The 4,4-MDA moiety was stabilized through hydrophobic interactions with Trp130 from below, and with Leu303 and Phe307 from above (Figure 7C), which together sandwiched the substrate’s benzene rings between their hydrophobic side chains. The flexible polyol chain attached to the target carbamate group (lodged in the oxyanion hole) was transiently stabilized by Thr212 and Val306 through hydrophobic contacts with its ethylene groups, and by Ser215, which donated an H-bond to the polyol chain’s oxygen atoms (Figure 7D). The second polyol chain, located at the entrance of the binding site, remained largely solvent-exposed and less tightly bound to the enzyme, which enabled it to explore a broader conformational space.

We also examined the binding dynamics of MUE-MDA within u15’s binding site. Interestingly, the docking results again clustered into two different poses, Pose A and Pose B, as observed for DUE-MDA (Figure S8A). MD simulations consistently revealed that Pose A was more stable and catalytically competent than Pose B (Table S2). All key interactions identified for DUE-MDA were preserved in the u15:MUE-MDA complex, with both the 4,4-MDA moiety and the polyol chain linked to the target urethane bond occupying the same regions as their counterparts in the u15:DUE-MDA complex (Figure S8B), although it is not a kinetically favorable substrate.

### Hydrolysis of Flexible PUR Foam

To evaluate the hydrolytic potential of the enzyme panel on industrially relevant polyurethanes, we first performed a proof-of-concept assay using a diaminotoluene (TDA) based polyurethane fragment, DUE-TDA (Figure S1B). This simplified substrate enabled real-time monitoring of monomer release, specifically 2,4-diaminotoluene (2,4-TDA), providing an initial indication of enzymatic cleavage efficiency. The assay was also valuable for comparing enzyme activity on TDA- versus MDA-based polyurethanes, helping to identify differences in substrate recognition prior to testing on the more complex industrial material. The half-hydrolyzed intermediates monourethane-2-diaminotolune, MUE-2-TDA, and monourethane-4-diaminotolune, MUE-4-TDA, were detected as the predominant product (Figure S1B); however, in the absence of a suitable standards, only DUE-TDA and monomeric 2,4-TDA were quantified (Figure S9). Within two hours, u17 released 0.2 mM of 2,4-TDA, followed by u15 producing 0.1 mM (Figure S9A, Figure S9B). The candidate u16 reached a plateau after one hour, releasing 30 µM 2,4-TDA, while the reference enzyme SP1 showed comparable activity to u13, both reaching approximately 30 µM 2,4-TDA after 1.5 hours (Figure S9B). Building on these preliminary results, all urethanases (u10, u13, u15-17) and SP1 were subsequently tested on a generic flexible PU foam (Figure S1C), derived from 2,4- and 2,6-toluene diisocyanate (TDI).^29^ After 24 h, u13, u15, u17, and SP1 emerged as the most active candidates, mainly releasing 2,4-TDA at concentrations of 2.5 µM, 2.7 µM, 2.5 µM, and 1.9 µM. After 48 h, the 2,4-TDA concentration increased to 9.7 µM for u13, 10.0 µM for u15, 9.2 µM for u17, and 6.1 µM for SP1. An increase in 2,6-diaminotoluene (TDA) was also detected after 48 h, reaching 8.0 µM for u13, u15, and u17, and 3.0 µM for SP1 (Figure S10). These findings confirm that the urethanases can act on industrially complex substrates such as flexible foam, highlighting that the major determinants of enzymatic performance are substrate accessibility and enzyme stability under the reaction conditions. Both parameters are critical targets for optimization to achieve more efficient hydrolysis, for instance by combining complementary enzymes in synergistic reaction setup.

### Conclusions

This study reports the discovery and analysis of five new and unique bacterial urethanases identified through bioinformatic data mining with the reference enzymes UMG-SP1-3 ^28^. Their activities toward polyurethane-like substrate were evaluated using the model compound DUE-MDA, which was cleaved by all variants to yield the mono-urethane intermediate MUE-MDA. Kinetic characterization revealed significant differences in catalytic performance, with u15 displaying the highest turnover rate (k_cat_ = 14.6 s⁻¹) and catalytic efficiency, outperforming with3-fold increase the reference enzyme SP1. Time-course analyses confirmed the conversion of DUE-MDA to MUE-MDA across all variants, with u15 and u17 achieving detectable further hydrolysis of MUE-MDA to release 4,4-MDA, highlighting variant-specific differences in catalytic specificity. Structural analysis of u15 provided the molecular basis for its high activity. The crystal structure, determined at 2.05 Å in complex with ethylene glycol, revealed a compact α/β-fold characteristic of the amidase signature (AS) family, containing the canonical Ser177–cis-Ser153–Lys78 catalytic triad supported by Ser154 and Ser172. The active site pocket, predominantly hydrophobic, guides substrates toward the catalytic center, with key residues including Leu303, Phe307, Trp130, and Gln365 shaping its geometry. Comparative analysis with the closely related enzyme SP3 revealed local variations in loops L3 and L4, as well as specific residue substitutions that likely modulate substrate binding and product release. Complementary docking and molecular dynamics simulations corroborated these structural observations, revealing that only one substrate orientation (Pose A) (Figure S6A) maintained catalytically favorable alignment, stabilizing the carbamate group within the oxyanion hole and enabling efficient nucleophilic attack by Ser177. The combination of these structural and dynamic features explains the high catalytic competence of u15 across urethane substrates. To assess performance on industrially relevant TDI-based substrates, the enzyme panel was tested on the soluble model compound DUE-TDA and a representative flexible foam substrate, where u15 and u17 confirmed to be most active candidates. Altogether, this work not only expands the diversity of characterized urethanases but also identifies structural and mechanistic determinants that govern activity and substrate adaptation. These insights lay the groundwork for engineering urethanases with enhanced efficiency, substrate flexibility, and stability, accelerating the development of enzymatic tools for sustainable polyurethane recycling and the broader mitigation of plastic waste.

## ASSOCIATED CONTENT

### Supporting Information

The following files are available free of charge. Hydrolysis reaction on DUE-MDA, DUE-TDA and Flexible PUR foam; Screening assay using Chromophore; HPLC chromatograms and calibration curves; Data collection and refinement statistics; Docking solutions for DUE-MDA; Distributions of the interatomic distances for Pose A and Pose B; Docking solutions for MUE-MDA; Average key interatomic distances and corresponding standard deviations obtained from the simulations of the u15:DUE-MDA and u15:MUE-MDA complexes; Progress curves on DUE-TDA and relative standard courves; Hydrolysis of Flexible PUR Foam.

## ACKNOWLEDGMENT

EnZync is supported by Challenge grant NNF22OC0072891 from the Novo Nordisk Foundation. We are grateful to the rest of the EnZync consortium members for constructive and helpful feedback. PP, PAF, and MJR acknowledge the financial support from the PT national funds (FCT/MECI, Fundação para a Ciência e Tecnologia, and Ministério da Educação, Ciência e Inovação) through the project UID/50006/2025 - Laboratório Associado para a Química Verde - Tecnologias e Processos Limpos. PP, PAF, and MJR would further like to thank the European High-Performance Computing Joint Undertaking (EuroHPC JU), which granted access to Deucalion, the petascale EuroHPC supercomputer located in Guimarães, Portugal, under proposal 2025.00056.CPCA.A3. We acknowledge DESY (Hamburg, Germany), a member of the Helmholtz Association HGF, for the provision of experimental facilities. Parts of this research were carried out at PETRAIII, and we would like to thank Kirill Kovalev for his assistance. Beamtime was allocated for proposal MX1043.

## ABBREVIATIONS

AS,: amidase signature
PUR,: polyurethane
DUE-MDA,: di-urethane ethylene Methylenedianiline
4,4-MDA,: 4,4-methylenedianiline
DUE-TDA,: di-urethane ethylene Diaminotoluene
2,4-TDA,: 

## TOC

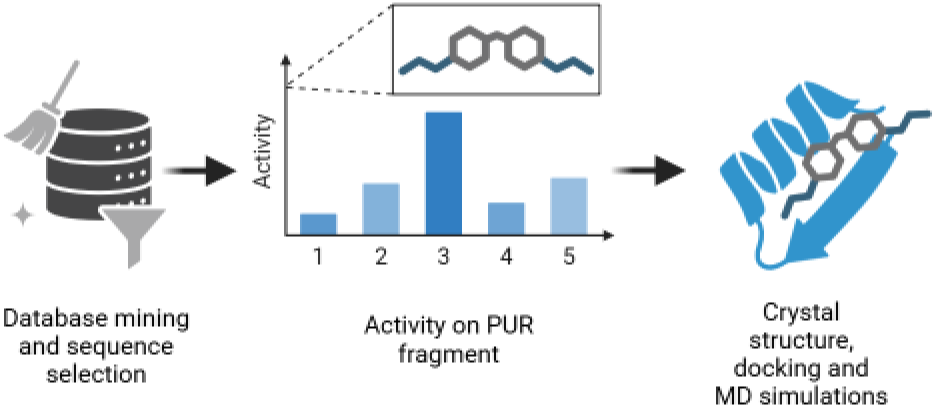

