## Supporting Information for "Expanding the Enzymatic Landscape for Polyurethane Degradation of Novel Bacterial Urethanases"

[b] Biotechnology and Biomedicine

Technical University of Denmark

Søltofts Plads, DK-2800, Kongens Lyngby, Denmark.

[c] LAQV, REQUIMTE, Departamento de Química e Bioquímica

Faculdade de Ciências, Universidade do Porto

Rua do Campo Alegre s/n, 4169-007 Porto, Portugal

[d] Danish Technological Institute

Kongsvang Alle 29, 8000 Aarhus, Denmark.

[e] Interdisciplinary Nanoscience Center (iNANO).

Aarhus University

Gustav Wieds Vej 14, 8000 Aarhus, Denmark

**Corresponding Author**

KEYWORDS amidase signature family• polyurethane degradation • enzyme catalysis • molecular docking • thermoset plastic • enzyme engineering

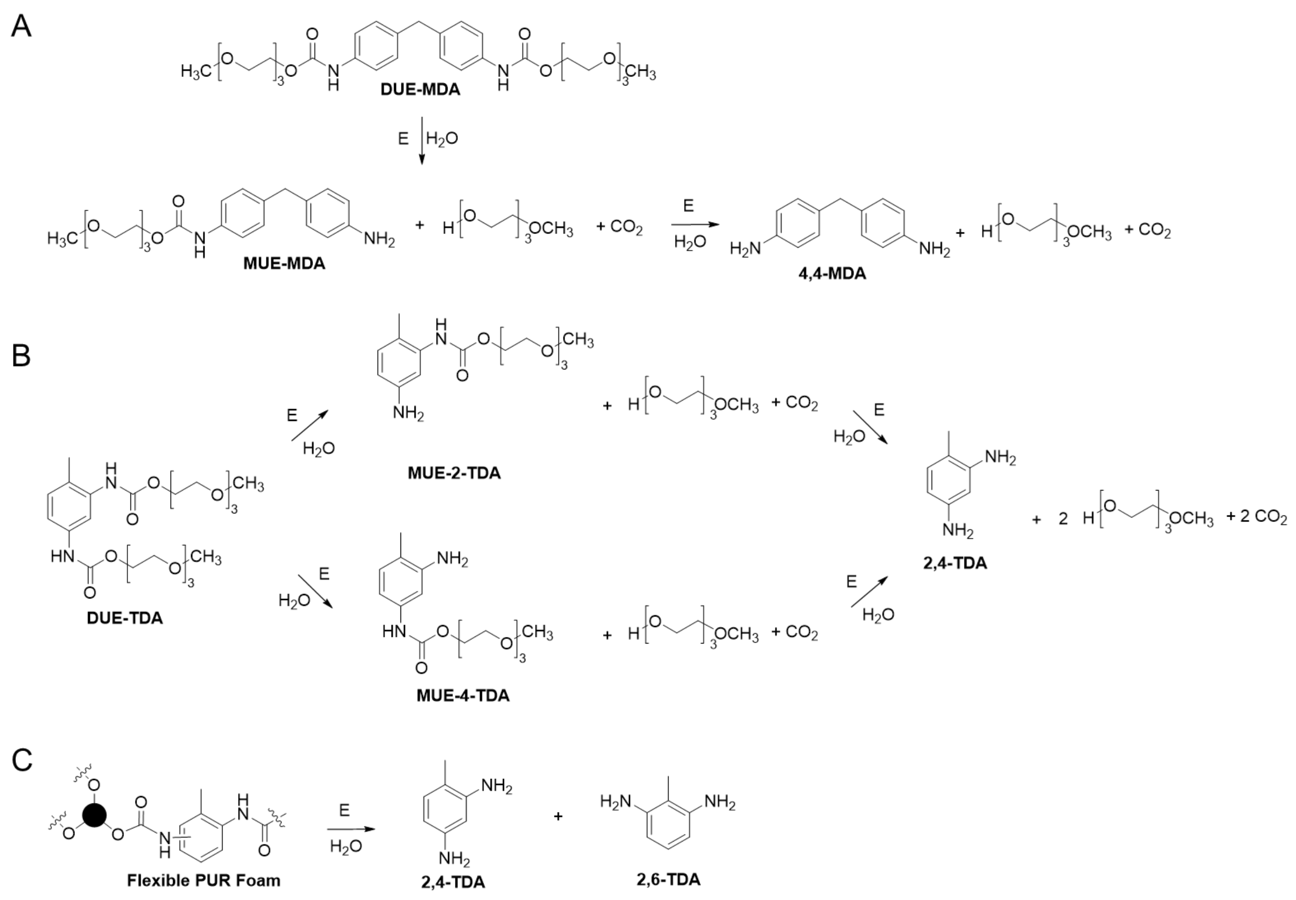


**Figure S1.** A) Hydrolysis reaction of DUE-MDA. Cleavage of di-urethane ethylene methylenedianiline (DUE-MDA) yields the monomeric 4,4-methylenedianiline, 4,4-MDA, the asymmetric monosubstituted mono-urethane ethylene Methylenedianiline, MUE-MDA, the 2-(2-(2-methoxyethoxy)ethoxy)ethan-1-ol sobstituent and carbon dioxide. B) Hydrolysis reaction of di-urethane diaminotoluene, DUE-TDA, producing two intermediate products monourethane-2-diaminotolune, MUE-2-TDA, and monourethane-4-diaminotolune, MUE-4-TDA, and the monomeric 2,4-diaminotoluene, 2,4-TDA, the 2-(2-(2-methoxyethoxy)ethoxy)ethan-1-ol substituent and carbon dioxide. C) Flexible PUR foam enzymatic hydrolysis yielding the two monomer 2,4-diaminotoluene, 2,4-TDA, and 2,6-diaminotoluene, 2,6-TDA. Structure of Flexible PUR Foam adapted from [1].

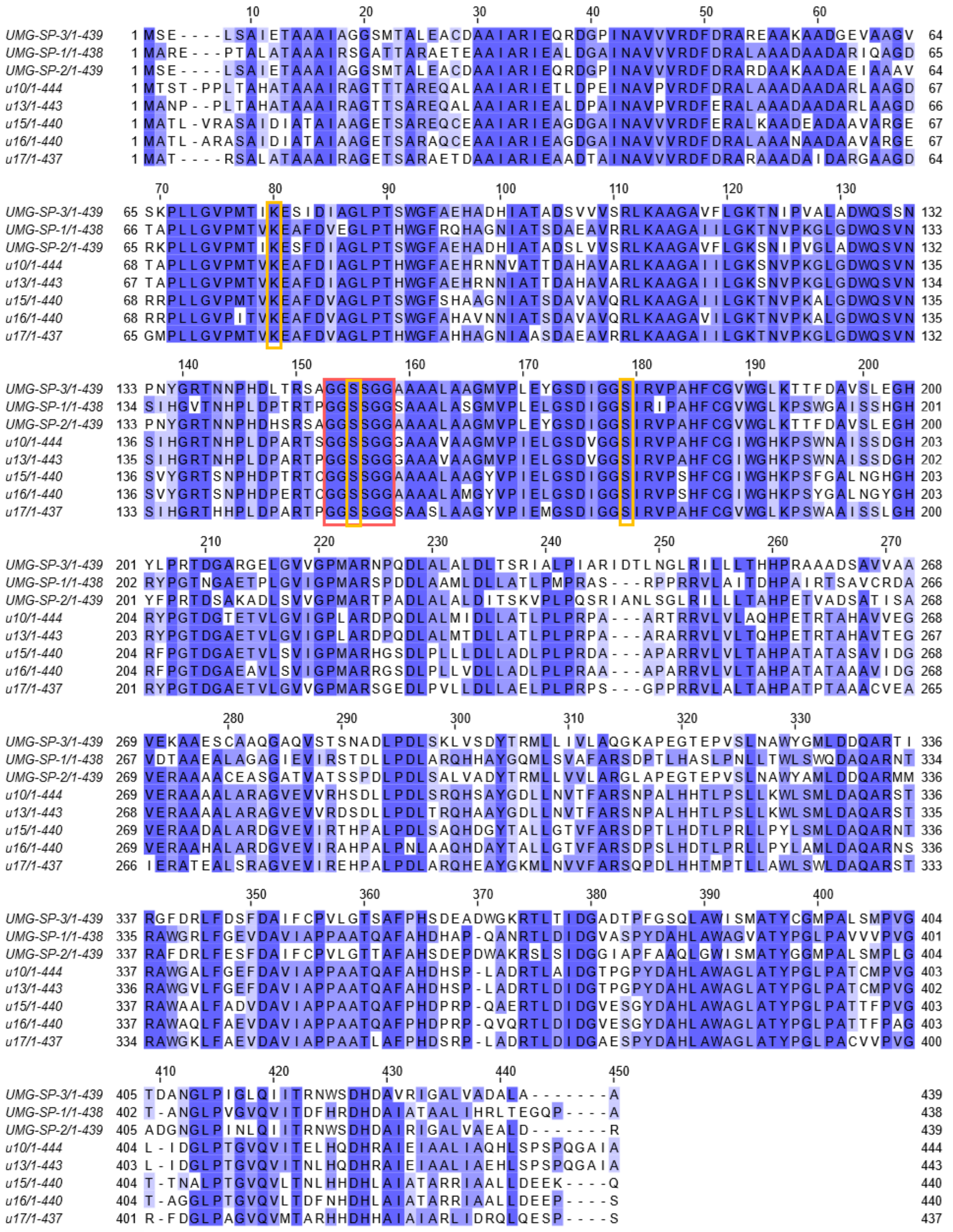


**Figure S2.** Multiple sequence alignment of the selected candidates u10-17 and the reference enzymes SP1-3. The sequences are colored by percentage of identity with 0 being 0% identical and 1 being 100% identical. Residues marked with yellow square represent the catalytic triad, residues in the red square represent the amidase signature motif.

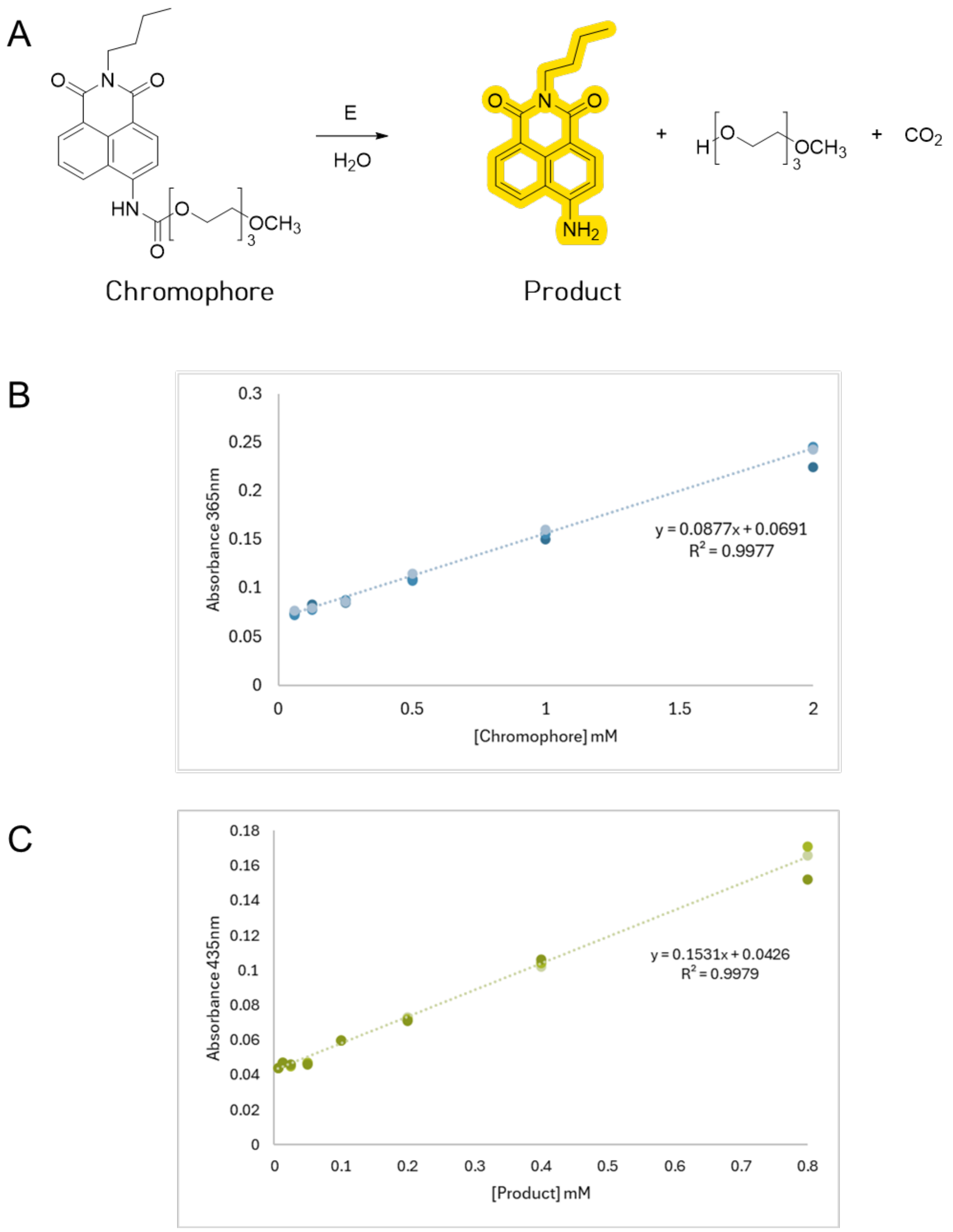


**Figure S3.** A) Screening assay based on Chromogenic substrate hydrolysis. The reaction monitors the product release over time at 435nm. B) Standard curve of chromogenic substrate measured at 365nm within the concentration range of 0.062-2 mM. B) Standard curve of the reaction product measured at 435nm within the concentration range of 0.015-0.8 mM.

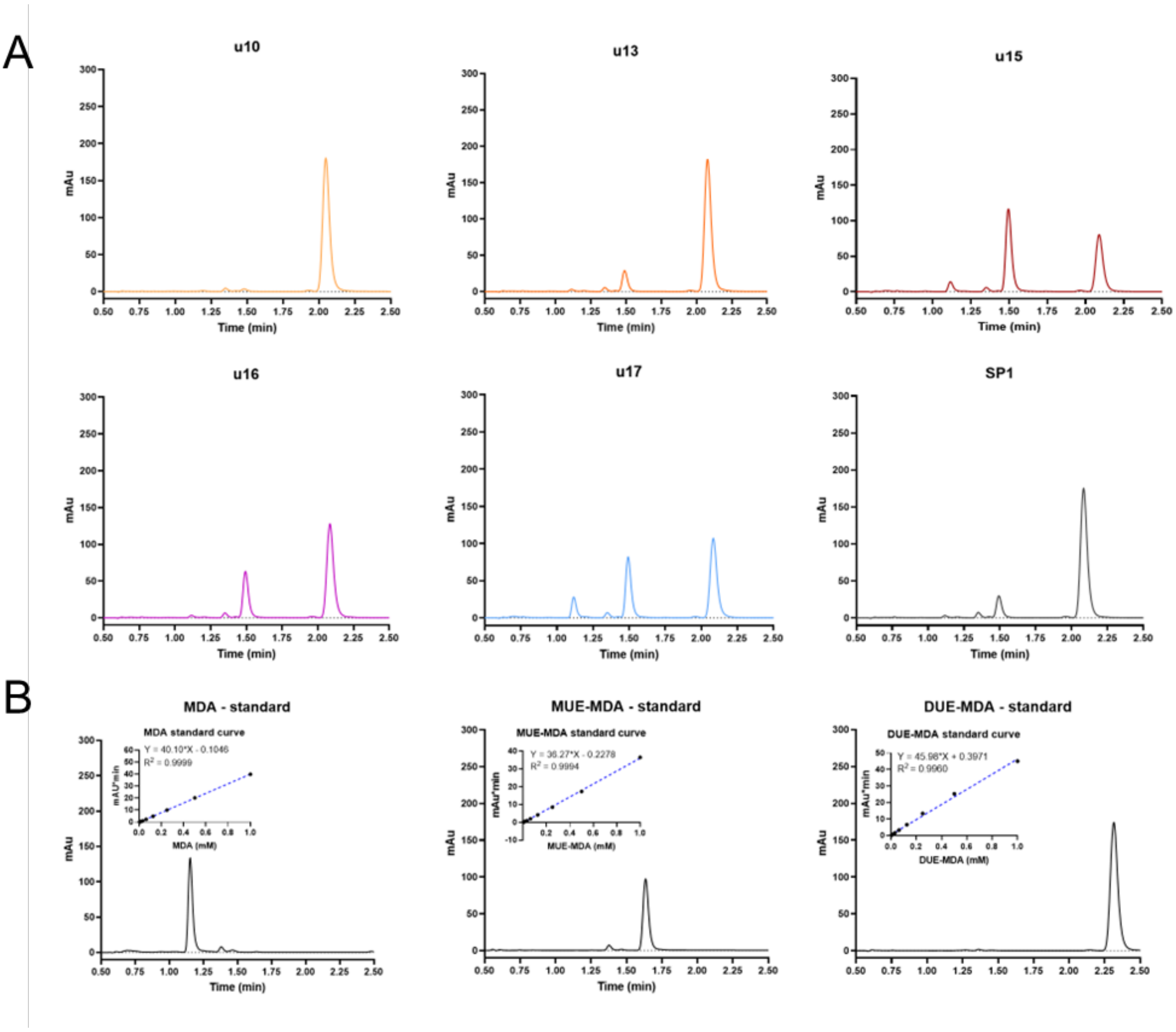


**Figure S4** A) HPLC Chromatograms of u10-17 and SP1 reaction mixtures B) Close-up view on MDA, MUE-MDA and DUE-MDA retention time and Calibration curves were interpolated with linear regression. Mean and standard deviations from two measurements.

**Table S1. Data collection and refinement statistics for u15 structure[a].**

| Structure | u15 |
|---|---|
| PDB ID | 9SK0 |
| Resolution [Å] | 46.45 - 2.052<br>(2.09 - 2.05) |
| Space group | P 21 21 2 |
| a. b. c [Å] | 95.956 106.154 86.286 |
| α. β. γ [°] | 90 90 90 |
| Total reflections | 604128 (504) |
| Unique reflections | 47045 (106) |
| $R_{merge}$ | 0.1632 (1.437) |
| $R_{meas}$ | 0.17 (1.606) |
| $R_{pim}$ | 0.04683 (0.6817) |
| CC½ | 0.994 (0.158) |
| I/σ (I) | 15.86 (1.25) |
| Completeness [%] | 96.34 (77.03) |
| Reflections for refinement | 53788 (2237) |
| Reflections for $R_{free}$ | 2615 (106) |
| $R_{work}$ | 0.1783 |
| $R_{free}$ | 0.2328 |
| No. of refined non-H atoms | |
| Protein | 6405 |
| Solvent | 515 |
| Ligands | 4 |
| Ramachandran Favored [%] | 97.36 |
| Ramachandran Allowed [%] | 2.64 |
| Ramachandran Outliers [%] | 0.00 |
| Rotamer outliers [%] | 1.41 |

[a] Values in parenthesis are for reflections in the highest resolution shell

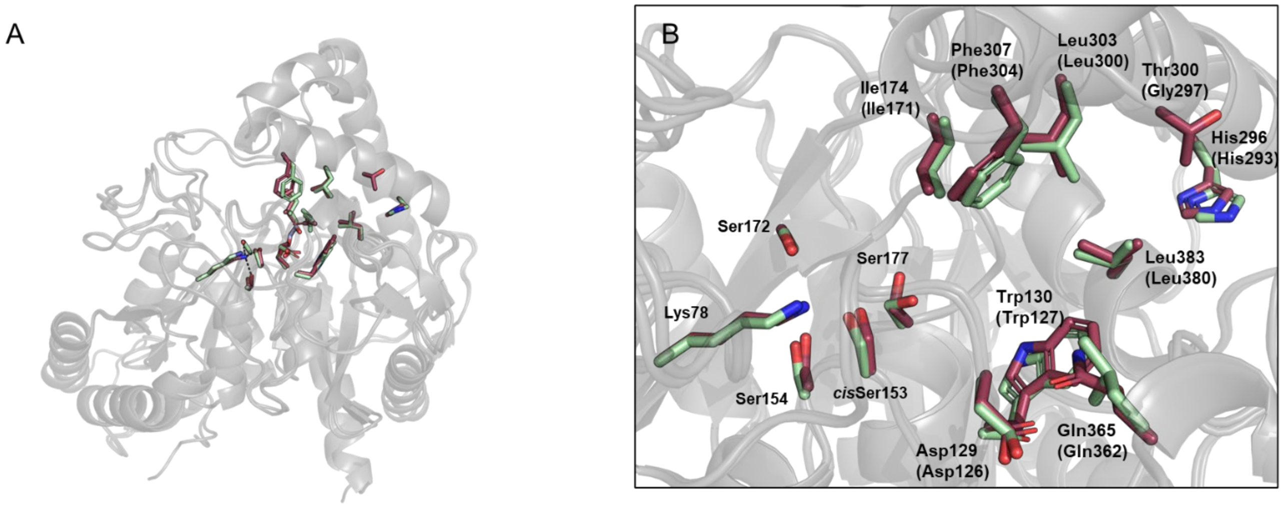


**Figure S5.** A) Comparison of the overall structure of u15 (PDB 9SK0 coloured in bordeaux) and the alpha fold model of u17 (coloured in light green). B) Differences in the binding pocket of u15 (coloured in bordeaux) and u17 (coloured in light green). Residues in brackets belong to u17.

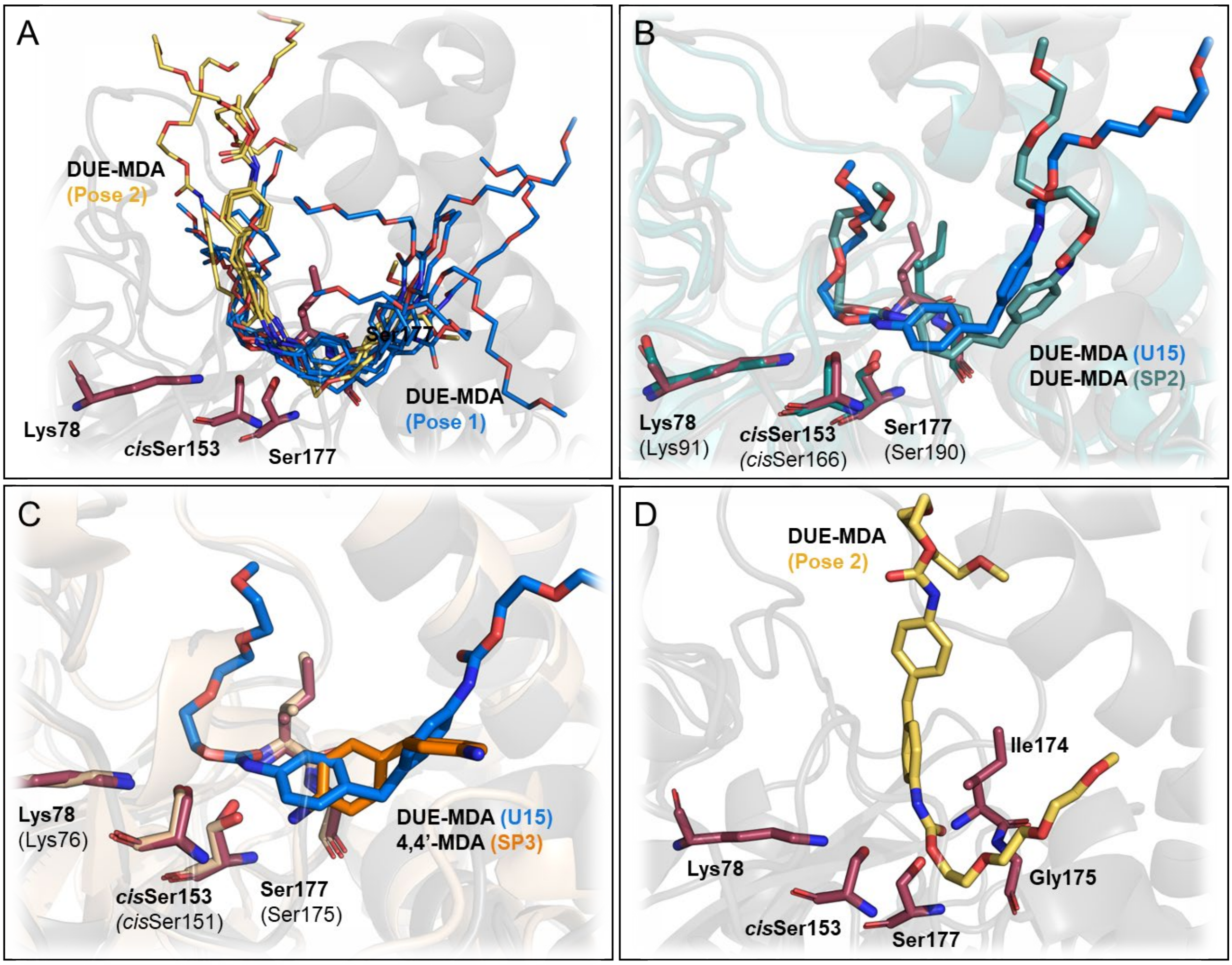


**Figure S6.** A) Docking solutions obtained for DUE-MDA within the u15 binding site. Solutions adopting Pose 1 are shown in blue, while those adopting Pose 2 are shown in yellow. B) Structural alignment of u15:DUE-MDA (best-scoring Pose 1) with UMG-SP2:DUE-MDA (complex used in previous computational studies), showing that Pose 1 closely resembles the configuration adopted by DUE-MDA in UMG-SP2. Residues in brackets correspond to UMG-SP2. C) Structural alignment of u15:DUE-MDA (Pose 1) and UMG-SP3:4,4'-MDA (PDB ID: 9FZ1) complexes, showing that the 4,4'-MDA moieties occupy the same spatial region. Residues in brackets correspond to UMG-SP3. D) Best-scoring docking result for DUE-MDA in Pose 2.

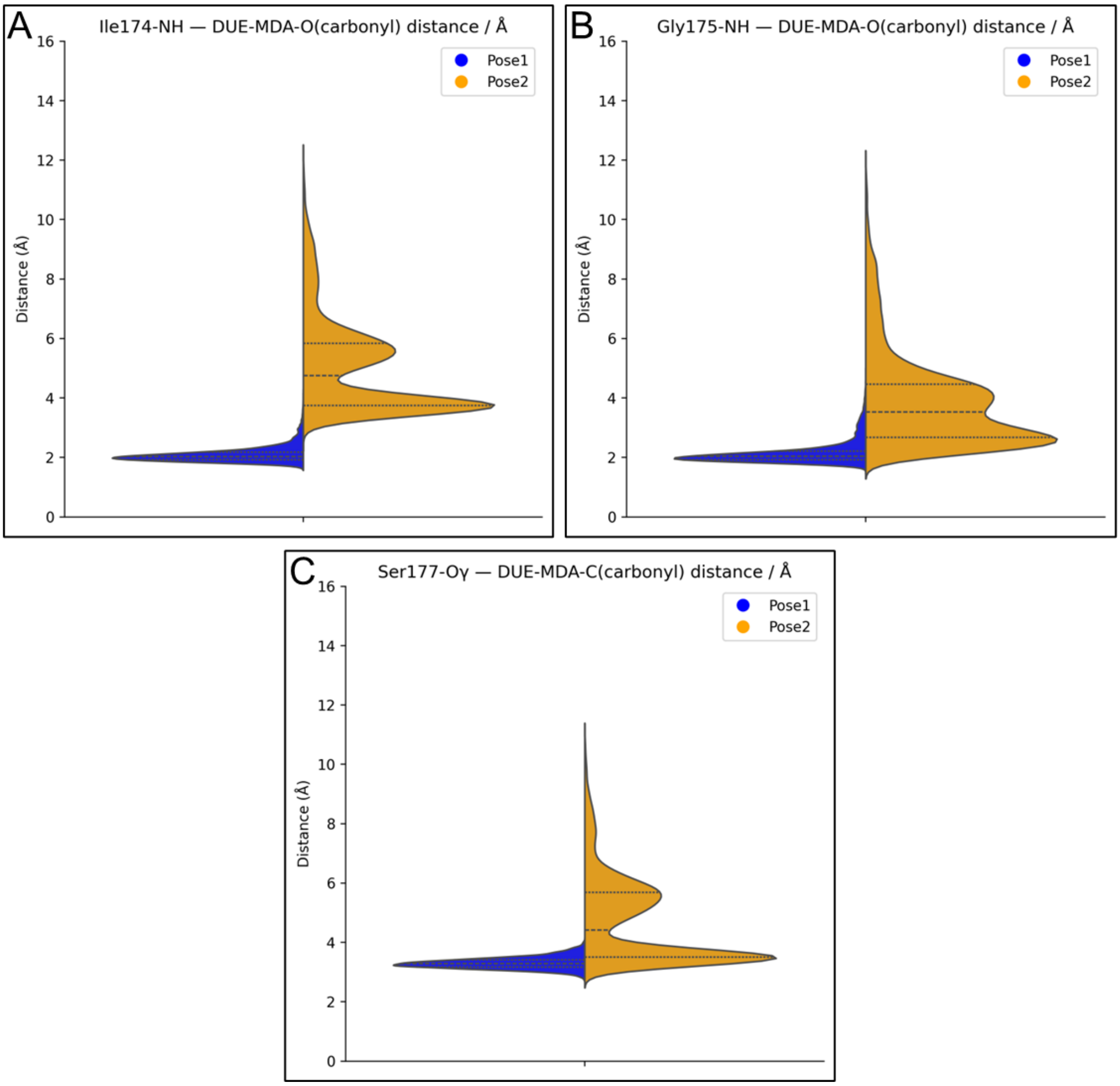


**Figure S7.** Distributions of interatomic distances for substrate Poses 1 and 2 between the DUE-MDA carbonyl oxygen and the backbone amides of (A) Ile174 and (B) Gly175, and between the DUE-MDA carbonyl carbon and the Oγ atom of Ser177 (C).

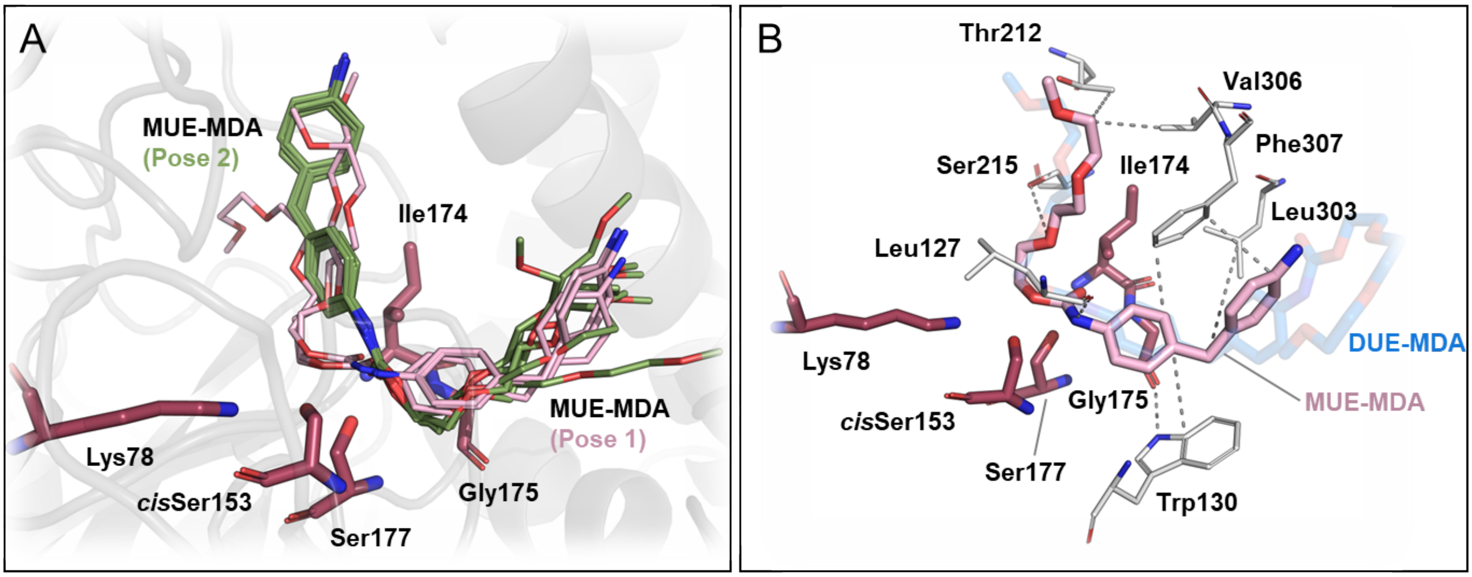


**Figure S8.** A) Docking solutions obtained for MUE-MDA within the u15 binding site. Solutions adopting Pose 1 are shown in bright pink, while those adopting Pose 2 are shown in green. B) Key interactions within the u15:MUE-MDA (best-scoring Pose 1) docking complex, highlighting residues involved in substrate binding. The DUE-MDA Pose 1 complex is overlaid in transparent blue sticks for comparison, illustrating the close alignment between both substrates.

**Table S2.** Average key interatomic distances and corresponding standard deviations obtained from the 1.2 μs MD simulations of the u15:DUE-MDA and u15:MUE-MDA complexes.

| | **Interatomic distance** | **Pose 1** | **Pose 2** |
|---|---|---|---|
| **DUE-MDA** | $O_{carbonyl}$(DUE-MDA) – NH(Ile174) | 2.1 ± 0.3 Å | 5.1 ± 1.6 Å |
| | $O_{carbonyl}$(DUE-MDA) – NH(Gly175) | 2.1 ± 0.4 Å | 3.9 ± 1.6 Å |
| | Nζ(Lys78) – Hγ(*cis*Ser153) | 1.8 ± 0.1 Å | 1.8 ± 0.2 Å |
| | Oγ(*cis*Ser153) – Hγ(Ser177) | 2.9 ± 0.6 Å | 3.9 ± 0.3 Å |
| | Oγ(Ser177) – $C_{carbonyl}$ (DUE-MDA) | 3.3 ± 0.2 Å | 4.7 ± 1.5 Å |
| **MUE-MDA** | $O_{carbonyl}$(MUE-MDA) – NH(Ile174) | 2.3 ± 0.5 Å | 7.8 ± 3.4 Å |
| | $O_{carbonyl}$(MUE-MDA) – NH(Gly175) | 2.4 ± 0.7 Å | 6.3 ± 3.4 Å |
| | Nζ(Lys78) – Hγ(*cis*Ser153) | 2.0 ± 0.5 Å | 1.8 ± 0.4 Å |
| | Oγ(*cis*Ser153) – Hγ(Ser177) | 3.0 ± 0.7 Å | 3.8 ± 0.3 Å |
| | Oγ(Ser177) – $C_{carbonyl}$ (MUE-MDA) | 3.4 ± 0.3 Å | 7.1 ± 3.3 Å |

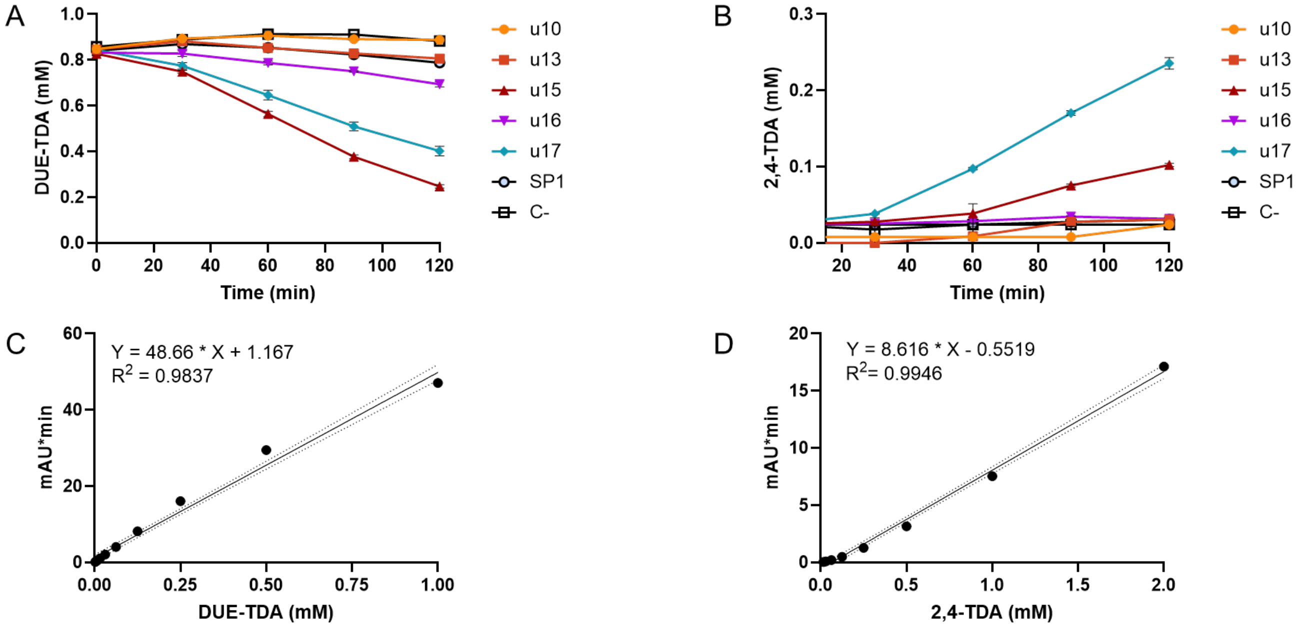


**Figure S9.** A) Hydrolysis reaction performed on DUE-TDA monitored over time. B) Release of 2,4-TDA over time. C) Standard curve of DUE-TDA D) Standard curve of 2,4-TDA. The standard curves are interpolated with linear regression. Mean and standard deviations from three measurements. Reactions contained 0.1 μM enzyme and 1 mM DUE-TDA (maximum 5% ethanol, v/v) in 50 mM sodium phosphate buffer, pH 8.0, and were performed at 40 °C with shaking at 800 rpm. The injection volume was 5 μL.

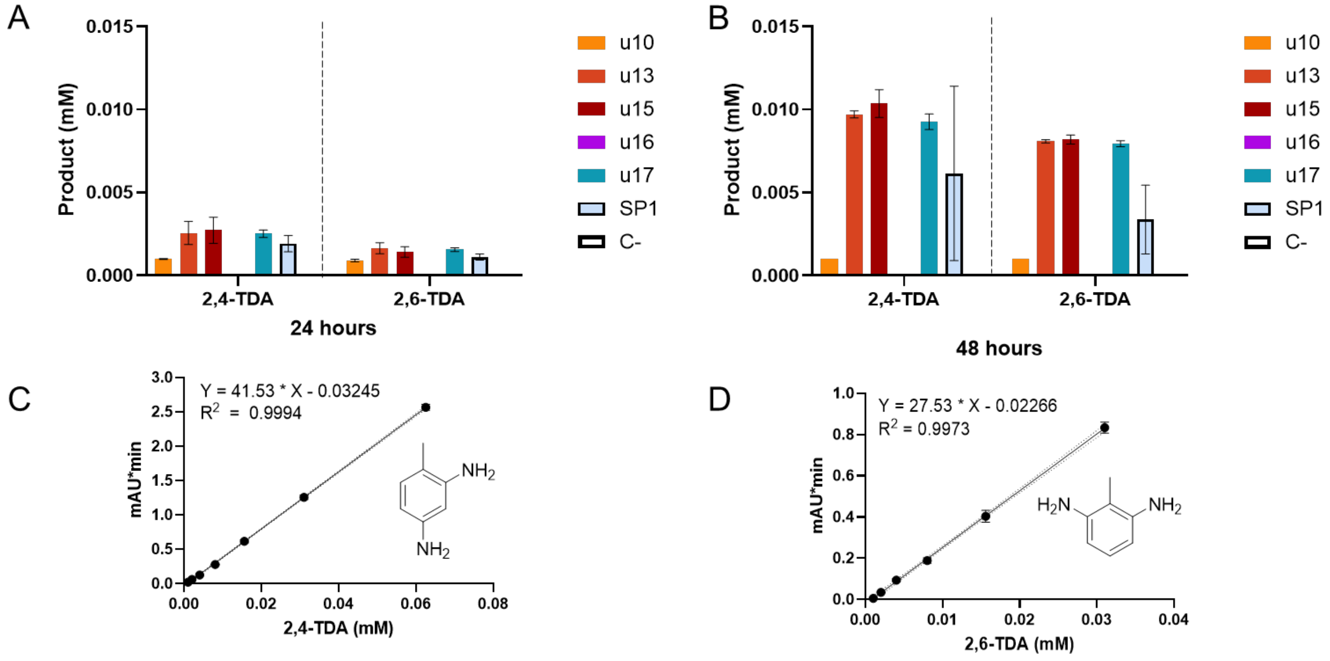


**Figure S10.** A) Quantification of monomeric 2,4-TDA and 2,6-TDA after 24 hours of incubation. B) Quantification of monomeric 2,4-TDA and 2,6-TDA after 48 hours of incubation. C) Standard curve of 2,4-TDA. D) Standard curve of 2,6-TDA. The standard curves are interpolated with linear regression. Mean and standard deviations from three measurements. The reaction consists of 5µM enzyme and 25 mg/ml Flexible Foam (maximum 5% Ethanol (v/v)) in 50 mM Sodium Phosphate buffer (pH 8). at 40 °C with 800 rpm shaking. The injection volume was 20 µL.
